# The E34 Phage Tailspike Protein: An in vitro characterization, Structure Prediction, Potential Interaction with *S. newington* LPS and Cytotoxicity Assessment to Animal Cell Line

**DOI:** 10.1101/2021.09.20.461090

**Authors:** Joseph A. Ayariga, Logan Gildea, Honghzuan Wu, Robert Villafane

**Affiliations:** The Biomedical Engineering Program, College of Science, Technology, Engineering and Mathematics (C-STEM), Alabama State University, 1627 Hall Street, Montgomery, AL, 36104; The Microbiology Program, College of Science, Technology, Engineering and Mathematics (C-STEM), Alabama State University, Montgomery, AL, 36104; Department of Biological Sciences, College of Science, Technology, Engineering and Mathematics (C-STEM), Alabama State University, 1627 Hall Street Montgomery, Alabama 36104; Department of Biological Sciences, Microbiology Ph.D. Program, College of Science, Technology, Engineering and Mathematics (C-STEM), Alabama State University, 1627 Hall Street Montgomery, Alabama 36104

## Abstract

The E34 phage is a member of the podoviridae family of phages, (short non-contractile tailed bacteriophages) that uses *Salmonella newington* as its host. This phage initiates the infection of its host via a specific interaction between its tailspike protein (TSP) and the lipopolysaccharides (LPS) of the bacterial. The E34 TSP is structurally similar and functionally equivalent to the P22 phage whose TSP has been well characterized and electron micrographs of both phages appear indistinguishable. The crystal structure of P22 phage TSP in complex with the O-antigen of *S. typhimurium* has been determined; and the active site of the TSP demonstrated to be the residues Asp392, Asp395 and Glu359 of the receptor binding domain. In another phage called E15, a phylogenetic relative of E34 phage, a short polysaccharide consisting of α repeating units is responsible for the interaction between the E15 phage and *Salmonella anatum’s* LPS leading to the adsorption of the phage to the bacteria. Studies on E34 phage shows that it interacts with *Salmonella newington’s* O antigen polysaccharide component of the LPS, this polysaccharide consists of mannosyl-rhamnosyl-galactose repeating units joined together by β-galactosyl linkages. However, no data exist regarding the specific residues of E34 TSP that are responsible for LPS binding and hydrolysis. In this study, the tailspike gene was cloned onto vector pET30a-LIC and expressed as a fusion protein termed the extended E34 TSP (EE34 TSP). We characterized the protein based on resistance to heat, SDS, and proteases; showing that the protein is heat resistant, shows aberrant electrophoretic mobility in the presence of SDS gradient, and actively binds to P22 phage heads to form hybrid phages that cannot infect P22 host. We also demonstrate via *in silico* study that the E34 TSP binds to and hydrolyses the O-antigen of its host via the ALA250, SER279 and ASP280 residues. Finally, testing E34 phage ability to protect Vero cells from Salmonella infection shows highly encouraging results, implying that E34 phage can be used in therapeutic/preventive medicine.

## 1. Introduction

Salmonellosis and typhoid are among the most frequently reported food and waterborne diseases worldwide with various transmission vehicles such as commercial chicken meat, eggs, vegetables, etc. In the United States alone, the Center for Disease Control (CDC) estimates that *Salmonella* bacteria cause over 1.35 million infections annually, with over 26,500 resulting in hospitalization, and nearly 420 deaths. Its symptoms include diarrhea, fever, and stomach cramps **(CDC)** ^1^. While most people can recover from it without treatment, antibiotic administration is used to treat people who show severe illness and are hospitalized. *S. newington* infections are infrequent and sporadic, unlike most common types such as *S. typhimurium* **(Neva et al., 1951)**.

Currently, antibiotic resistance is a significant threat to public health, in the United States alone, over 2.8 million people get antibiotic-resistant infections **(CDC^2^; Herhaus and Dikic, 2017)**. Fighting this threat demands a multidisciplinary and collaborative approach. One of the new methods of tackling antibiotic-resistant infections is the use of phages **(Alves-Barroco et al., 2020)**. Phages have extensive utility in science and engineering beyond disease control **(Zeng et al., 2018; Rehman et al., 2019; Silpe et al., 2018; Pausch et al., 2020).** Despite phages being one of the most abundant “life forms” on the planet **(Harada et al., 2018),** only a minuscule number of these bacterial viruses have been studied in any detail. Majority of phage utility come from the use of the phage’s tailspike protein which exhibits specificity toward their host receptors **(Zeng et al., 2018)**. *Salmonella* and their phages are a prototypical infection pair. The phages in many of these pairs recognize the lipopolysaccharide (LPS) moiety as their initial cellular receptor **(Lu et al., 2020)**. The tail proteins are a very important class of phage proteins that often determine the host specificity and are critical elements in the phage biology, especially the phage’s assembly pathway prior to cell lysis as well as host receptor attachment, binding, and phage entry **(Jäckel et al., 2015)**. Phage tails have been weaponized as therapeutic agents against specific sets of bacterial strains, especially pathogenic bacteria **(Al-Zubidi et al., 2019)**. There are a very large number of applications of phage tails **(Schmidt et al., 2016; Handa et al., 2008;** Jung et al., 2017; Kelly et al., 2015; Kunstmann et al., 2018; Marti et al., 2013; Philippe and Rodolphe, 2010; Schmelcher and Loessner, 2014; Nilsson et al., 2000).

The E34 phage is a podovirus that infects *Salmonella newington* **(Greenberg et al., 1995)**. Electron micrographs of both phages appear indistinguishable **(Parent et al., 2012)**. The crystal structure of P22 phage TSP in complex with the O-antigen of *S. typhimurium* has been determined; and the active site of the TSP demonstrated to be the residues Asp392, Asp395 and Glu359 of the receptor binding domain **(Stienbacher et al., 1997)**. In another phage called E15, a phylogenetic relative of E34 phage, a short polysaccharide consisting of α repeating units is responsible for the interaction between the E15 phage and *Salmonella anatum’s* LPS **(Takeda and Uetake, 1973)** leading to the adsorption of the phage to the bacteria. Studies on E34 phage shows that it interacts with *Salmonella newington’s* O antigen polysaccharide component of the LPS, this polysaccharide consists of mannosyl-rhamnosyl-β-galactosyl linkages **(Iwashita and Kanegasaki, 1975)**. However, no data exist regarding the specific residues of E34 TSP that are responsible for LPS binding and hydrolysis.

The P22 phage TSP possesses unusual thermal stability, protease resistance, and SDS resistance **(Greenberg et al., 1995)**. The 216 kDa TSP of the P22 phage is well characterized **(Chen & King, 1991; Steinbacher et al., 1994)** and its crystal structure consists of a dome-shaped N-terminal domain, a solenoid-shape central parallel beta-helix domain that is required in LPS binding, and a beta-prism domain that is associated with the function of clamping and stabilizing the trimeric protein **(Steinbacher et al., 1997)**. Even though there exists a significant homology between the E34 phage TSP and the P22 phage TSP, these TSPs used for attachment and hydrolysis of their LPS are not interchangeable, hence E34 phage incubation with *S. typhimurium* (the bacterial host of P22 phage) does not result in binding of E34 TSP to the LPS of *S. typhimurium* **(Salgado et al., 2004; Zayas & Villafane, 2007)**.

LPS is a major receptor for many phages, including the two phages that are the primary focus of this report (P22 and E34 phages) **(Broeker and Barbirz, 2017; Ayariga et al., 2018; Kunstmann et al., 2018; Casjens and Hendrix, 2015)**. In many podoviruses DNA injection is preceded by the hydrolysis of the LPS of their respective hosts before anchoring the phage particle to the cell surface **(Broeker and Barbirz, 2017; Koskella and Brockhurst, 2014; Salgado et al., 2004)**. In the P22 infection process, phage P22 TSP binds and hydrolyses its host’s LPS in a coupled fashion as the phage works its way down to the bacterial cell surface **(Broeker & Barbirz, 2017; Ayariga et al., 2018)**. This is followed by the subsequent orientation of the viral particle on the cell surface for infection of the *Salmonella* host. Phage proteins located within the phage are released upon infection and are involved in the formation of a transmembrane structure **(Ayariga et al., 2018; Andres et al., 2012; Steinbacher et al., 1994)**. The E34 phage is proposed to follow a similar infection process **(Villafane et al., 2008)**. In the E34 phage system, most of the studies have been limited to structural homology studies especially between (E34 and P22-like phages) **(Salgado et al., 2004; Zayas and Villafane, 2007)**. Currently, there exist no in-depth study and characterization of the E34 phage components. E34 phage TSP, a 196.5 kDa trimer, is predicted to share similar structural topology with P22 TSP, with the head binding domains of both proteins sharing approximately 70% identity and confirmed to equally bind the capsid of P22 phage **(Zayas and Villafane, 2007)**.

In this work, we created the EE34 TSP by adding at the N-terminal domain of the protein a 43 amino acid fragment containing a His-tag sequence during the cloning process. This enabled our characterization of the properties of the EE34 TSP. Hence, the stability of this new protein regarding heat, SDS treatment, and the combination of the two have been examined. Also, the protein’s activity has been tested. This study reveals significant differences that distinguish the E34 TSP from the well-studied P22 TSP. Furthermore, we offer a predictive analysis of the possible interaction of the E34 TSP and its ligand (the O-antigen of S. newington LPS) using *in silico* docking studies.

There is an increased antibiotic resistance, this has caused a renewed interest in the use of phages to control and treat bacterial pathogens **(Meader et al., 2010; McVay et al., 2007; Yosef et al., 2014; Bragg et al., 2014)**. Hence evaluating the safety and efficacy of phages used in phage therapy is crucial. Wall et al., 2010 showed that the administration of phage cocktail to pigs contaminated with *Salmonella* typhimurium, reduce cecal *Salmonella* concentrations by 95% and ileal *Salmonella* concentrations by 90%. They demonstrated that the phage cocktail was lytic to several non-*Typhimurium* serovars **(Wall et al., 2010)**. In a similar study, direct feeding of microencapsulated bacteriophages to pigs reduces salmonella colonization **(Saez et al., 2011)**. In this work, Saez et al., 2011 fed pigs with bacteriophage with microencapsulated bacteriophages, and demonstrated that the feed group were less likely to shed *Salmonella* typhimurium at 2uh: 71.4%; 4uh: 71.4%; 4 h: 85.7%) groups (*p*<0.05) **(Saez et al., 2011)**. Another method of assessing the efficacy of phages is the employment of *in vitro* studies, which are crucial to understanding the complex and dynamic interactions between phages and animal cells.

Similar to drug screening, animal cells have been employed to screen the efficacy of several phages and phage cocktails **(Shan et al., 2018; Nielsen et al., 2002; Abedon et al., 2017; Shan et al., 2012)** to demonstrate the significance of cell-line in assessing safety of phages. However, no literature exists that informs of the effects the E34 phage on animal cell lines we attempt an initial assessment of the phage’s cytotoxicity and its effects on Vero cells viability. Our future goal is to use the cloned and expressed tailspike of this phage to study its bacteriostatic property and to evaluate its ability to inhibit *S. newington* growth in *in vitro* cultures.

## 2. Materials and Methods

### 2.1. Media, chemicals, and other reagents

Kanamycin-resistant (KmR) transformants were grown on LB agar medium premixed with the antibiotic at 50 mg/ml. A total of 1 mM isopropyl-β-D-galactopyranoside (Sigma, New York, USA) was used for induction. SOB and SOC media were purchased from New England Biolabs (Newburyport, MA); also, Oligonucleotides, Taq polymerases, and other enzymes were from New England Biolabs. Competent Cells BL21/DE3 and Novablue cells were purchased from Novagen (Sigma Millipore, USA), P22 host cells; Salmonella *typhimurium* (BV4012) and BV7) came from our laboratory collection, *Salmonella newington*) was a kind gift from Sherwood Casjens (University of Utah). The pET30a-LIC vector was purchased from Novagen (Sigma Millipore, USA). Urea, NaCl, Ammonium sulfate, and all other chemicals used in this research were of HPLC grade.

### 2.2. pET30a-LIC-E34 TSP cloning and sequencing

E34 phage DNA was obtained as previously described **(Williams et al., 2019)**. The E34 TSP gene was copied from the complete E34 phage DNA via PCR using two primers (Forward and Reverse Primers, see Table 1) to generate our tailspike gene fragment. These primers have short special sequences at their ends to attach to vector LIC (ligation-independent cloning) ends. Using pET30a-LIC vector, which is a high expression vector that contains a T7 promoter, inducible by-D-1β thiogalactopyranoside (IPTG) and ligation independent. Therefore, it did not require the input of ligase. The pET30a-LIC vector adds an extra 43 amino acids which contain a His-tag in a short peptide fragment fused N-terminally to the TSP. The PCR reaction was performed using an Eppendorf S Master cycler programmed with the following temperature-cycling protocol: DNA denaturation at 95 °C for 2 mins, template denaturation at same 95 °C for 15 secs, primer annealing 55 °C for 10 secs, primer elongation 30 sec at 72°C, finally in its last cycle elongation of 5 mins at 72 °C. DNA denaturation, primer annealing, and primer elongation were repeated 35 times.

Putative clones were obtained after the LIC procedure. Clones containing the E34 TSP gene were verified by a PCR reaction using the primers listed in Table 1. The identity and veracity of the putative tailspike gene clones were obtained after DNA sequencing of the clones using Sanger sequencing at the Heflin Genomic Sequencing Center at the University of Alabama, Birmingham. The sequences obtained confirmed the sequence of the E34 TSP gene (Genebank Accession number DQ167568) **(Zayas and Villafane, 2007)**. The sequencing primers used were P1, P100, P312, and P504 (Table 1). The primers are named after the codon which their sequence contains, and they provide overlapping sequences that confirm published results when compared to the published sequence **(Zayas and Villafane, 2007)**. Plasmids were obtained from the verified clones and expressed. The expression protocol has been described **(Palmer et al., 2014, Sambrook et al., 1989)** and is briefly described below.

### 2.3. Extended E^34^ TSP expression and purification

A 5 mL of a freshly grown overnight cell of *Escherichia coli*; BL21/DE3 cells from Novagen containing the verified pET30a LIC-E34 TSP were diluted into a 1L flask containing 500 mL of LB broth with Kanamycin. The cells were then incubated at 37 °C with shaking in a MaxQ 4450 incubator (Thermo Scientific) with shaking until the culture reached an OD600 of 0.6 for induction. IPTG was added to a final concentration of 1 mM to induce cells. Growth after induction was carried out for 6 hours. Cells, after induction, were harvested by centrifugation and lysates, were generated as described previously **(Palmer et al., 2014)**. In brief, pelleted cells were then re-suspended in lysis buffer consisting of 50 mM Tris at pH 7.4, 5 mM MgCl2, 0.1 mg/mL lysozyme, 0.1 mg/mL DNase, 0.05 mg/mL RNASE, 0.2 mg/mL DTT and subjected to three cycles of freeze-thaw-freeze, and then finally centrifuged at 17,000 rpm for 30 mins. The supernatant was decanted into 50 mL tubes and stored at −20 °C as the EE34 TSP lysate. The target protein (EE34 TSP) was then fractionated using FPLC (GE/Amersham Biosciences-AKTA) connected to a desktop computer Pentium 4 running UNICORN software; fractions were pooled and enriched to the desired concentrations using Amicon concentrators (Millipore Sigma).

### 2.4. Recombinant enterokinase (rEK) digestion of EE^34^ TSP

To produce the matured TSP of E34 devoid of the extra 43-amino acid fusion peptide, we utilized rEK digestion to cleave off the fusion peptide enzymatically. Thus, the regular 606 amino acid E34 TSP product is obtained. The rEK enzyme recognizes and cleaves the specific sequence DDDDK located between the E34 TSP gene and the 43 amino acid tag. Digestion of extended E34TSP was done according to the recombinant Enterokinase user protocol TB150 Rev.C 0107(Novagen). The extent of proteolytic activity on our TSP was calibrated via SDS PAGE analysis.

### 2.5. SDS binding assay

The SDS binding assay was done as described by Ke Xia and his colleagues **(Ke et al., 2007)**; in brief, the EE34 TSP was incubated for 30 mins in different concentrations of SDS at pH 7.4. Aliquots of treated samples were taken and mixed with SDS-free loading buffer consisting of 50 mM Tris-HCl, 25 % glycerol, and 0.01 % bromophenol blue. Samples were then loaded without prior heating into a 12 % polyacrylamide electrophoresis (PAGE) gel well and run at 100 volts. The native (SDS-free) running buffer consisted of 25 mM Tris base and 0.2 M glycine. Gels were stained using Coomassie blue stain and imaged using Bio-Rad ChemiDoc, and densitometry was achieved via ImageJ software.

### 2.6. Thermal denaturation

The concentrations of protein samples were predetermined using the Quickdrop from Molecular Devices. The unfolding of our trimeric protein by heat was performed as described in **(Chen and King, 1991)** with few modifications. In brief, protein samples in 50 mM Tris-HCl, pH 7.4 were incubated at set temperatures (50 °C, 70 °C, 80 °C, and 90 °C), and at different time points, aliquots were withdrawn, and sample loading buffer consisting of 50 mM Tris-HCl, 2 % SDS, 5% 2-mercaptoethanol, 10 % glycerol, 0.03 % bromophenol blue, pH 7 were mixed and loaded into wells of 10 % SDS PAGE gels. Gels were fixed using a fixing solution consisting of methanol/acetic acid (50/10, v/v) then submerged into Coomassie blue-staining solution; next, the gels were destained using a destaining solution made of methanol/acetic (10/10, v/v). Gels were then photographed using the Bio-Rad ChemiDoc XRS+ programmed with the Quantity One software. The densitometric values were obtained using ImageJ software. Finally, the unfolding kinetics are illustrated via a graph plot and mean significance tested by one-way ANOVA and post-hoc comparisons of the means.

### 2.7. Spot assay for interference of native EE34 TSP on P22 phage assembly (qualitative study)

In this assay, samples of constant concentrations of P22 heads (0.078 mg/mL) were first incubated with EE34 TSP (0.48 mg/mL) for 30 mins, and the reaction was allowed at room temperature for the EE34 TSP to find and bind to the capsids of the P22 (termed heads of P22). Following this, varying concentrations of P22 TSP (2.0, 1.0, 0.5, 0.25, and 0.125 mg/mL) were added to the designated treatment mix and incubated at room temperature for another 30 mins. A negative control treatment was included in which P22 phage heads were incubated with EE34 TSP and then followed with buffer, while the positive control treatment consisting of an initial incubation of P22 heads with buffer (50 mM tris, pH 7) for the same time point, then counter titrated with P22 TSP. Another experiment included here was the use of extended P22 TSP (EP22 TSP) (a variant of P22 TSP that has the 43 amino acids cloned N-terminally to the protein similar to the EE34 TSP. A single drop (5 uL) of each treatment was spotted on an agar plate grown with *S. typhimurium;* plates were incubated at 37 °C overnight and then photographed.

### 2.8. Plaque assay for interference of EE34 TSP on P22 assembly (quantitative study)

20 mg/mL of EE^34^ TSP that was diluted serially (i.e., 20 mg/mL, 2 mg/mL, 0.2 mg/mL, 0.02 mg/mL, 0.002mg/ml, and 0.0002 mg/mL) and 200 ul of each dilution titrated with 200 uL of fixed concentration of P22 heads (0.078 mg/mL) in a ratio 1:1 v/v. The reaction was allowed to proceed under room temperature for 30 mins; afterward, 200 uL of a fixed concentration of P22 TSP (0.48 mg/mL) was added to the reaction mix and allowed to sit for an additional 30 mins.

200 uL of the reaction mixture was added to the top agar containing the host cells (*S. typhimurium*) and spread on an agar plate. The experiments were carried out in triplicates. Two control experiments were employed; the negative control had P22 heads treated with buffer and followed with EE34 TSP, whereas the positive controls treated P22 heads to buffer followed with P22 TSP. Plates were incubated at 37 °C overnight; then plaques were counted.

### 2.9. Protein sequence extraction for homology modeling of E34 TSP

The protein sequences of E34 TSP (Gene ID: 7353089) was extracted from the NCBI protein database^3^. The 3D structural modeling was performed by using Swiss-Modeler^4^, an online homology modeling and model evaluation program. Subsequently, the models’ quality and validation were assessed using structure assessment methods such as the QMEAN **(Benkert et al., 2011)** and Ramachandran plot analysis^5^.

### 2.10. Molecular Docking analysis of E34 TSP- the mannosyl-rhamnosyl-galactose (O-antigen derivative)

To investigate the interactions between the O-antigen of the LPS of the *S. newington* and the TSP of E34, we used an O-antigen substitute called the mannosyl-rhamnosyl-galactose (PubChem structure CID: 129729227) ligand, which is a derivative of the O-antigen of the *S. newington* LPS, the 2D structure of the ligand was downloaded from the PubChem database^6^ docked to E34 TSP using PyRx (an open-source software for performing virtual screening that combines AutoDock Vina, AutoDock 4.2, Mayavi, Open Babel, etc.). The modeled structure of E34 served as the receptor. The receptor was prepared using AutoDock Vina wizard, whereas the ligand was prepared using the Open Babel tool. In summary, the ligand structure was minimized and converted to a pdbqt format before uploading as a ligand. In the case of the receptor, the bond orders were assigned, and charged hydrogen atoms were added to the protein. The receptor structure was also minimized using the AutoDock Vina wizard. The protein was loaded into PyRx and converted to receptor; the receptor grid boxes were generated in PyRx using the built-in Vina Wizard module, grid boxes were maximized to cover all active sites of the receptors. Finally, AutoDocking of the ligand to the receptor was made using the AutoDock wizard in-built in PyRx program to blind dock the O-antigen moiety to the RBD of the E34 TSP with exhaustiveness of 9. This way, nine models were generated in the process, showing the interaction of the mannosyl-rhamnosyl-galactose with the E34 TSP at various binding sites. Each model was saved as a PDB file and exported into Biovia Discovery Studio software (version; 21.1.0.278) for specific atomic-atomic interaction analysis between the ligand and the receptor.

#### 2.10.1. Vero cell growth in varying concentrations of E34 phage

In this experiment, Vero cells were seeded at a density of 1 × 10^5^ into 96 well plates, then serial dilutions of E34 phages (2.33 × 10^2^ µg/ml to 2.33 × 10^-5^ µg/ml) were added to cells and incubated for 24 hours. The Vero cells-phage mixture was incubated at 37 °C, 5% CO2 for 24 hours. Wells were then washed twice with 1X PBS to remove dead cells in suspension and fixed with formaldehyde. Fixed cells were then permeabilized using 2% SDS solution and stained with trypan blue, washed twice again, and read at 600 nm in the Cytation 3 plate reader (Biotek, USA). As shown in Figure 19 above, the highest absorbance was recorded for cells treated with the lowest concentrations of E34 phages (2.33 × 10^-5^ µg/ml and 2.33 × 10^-5^ µg/ml µg/ml). An absorbance of 1.83 was recorded for E34 phage treatment at concentration of 2.33 × 10^2^ µg/ml, whereas, 2.29 was recorded for the lowest concentration of E34 phage treatment, thus producing a difference of 0.46 in absorbance between the two E34 phage concentration extremes. The highest absorbance (2.37) however was registered at 2.33 × 10^-4^ µg/ml.

#### 2.10.2. Proliferative inhibitory Effects of E34 phages on Vero cells using MTT assay

The effect of E34 phages on cell proliferation was determined by the MTT assays following similar procedure used by other researchers **(Cao et al., 2013)**. Briefly, Vero cells were cultured in high glucose DMEM supplemented with 10% FBS, 2 mM glutamine, 50 units/mL penicillin, and 50 mg/mL streptomycin. Cell proliferation assay was performed using the CellTiter 96^®^ AQ_ueous_ One Solution Cell Proliferation Assay (Promega). The phages were serially diluted and the appropriate aliquots added to adherent monolayer Vero cells growing in 96 well plates (cell density; 1.0 × 10^5^ cells/well). Administration of phages to cells were done concurrently with μL of 1X PBS (vehicle control) or the E34 phage solution was added into each well and incubated for 24 h. Subsequently, 20 μ well and incubated for 2 h, followed with reading at 492 nm using the Cytation 3 Biotek microplate reader. Cell proliferation inhibition was given by the expression: Fraction of inhibition = [1−(OD_cells+E34 phage_ – OD_blank_) / (OD_cells+PBS_ – OD_blank_)] where OD means the absorbance at 492 nm. Data presented as mean ± SEM, n = 3.

## 3. Results

### 3.1. Cloning and sequencing of tail-containing clone

The E34 gene was obtained after phenol extraction of a concentrated virus stock of the E34 phage as described **(Zayas and Villafane, 2007)**. The primers, as listed in Table 1, had a slight modification to work in the Ligation-independent system. The identity of the cloned E34 gene was obtained by agarose gel electrophoresis to indicate an increased molecular size of the plasmid (increased by the weight of the E34 gene) and reconfirmed by the DNA sequence of the cloned DNA by UAB Genomic Sequencing Center, Birmingham, Alabama (data not shown). The sequencing was done using the Sanger dideoxy methods using gene-specific primers P1, P100, P312, and P504 primers, Table 1. The sequencing data indicated sequence identity to the published data deposited in databases, Genebank Accession Number DQ167568 **(Villafane et al., 2008; Zayas and Villafane, 2007)**. The cloned tail gene was also verified by the removal of the fusion N-terminal addition of the fusion protein described below.

### 3.2. Extended E^34^ TSP expression and purification

The clone expressing the EE34 TSP as detected by SDS-PAGE analysis is seen in Fig.1B. It migrated at its predicted molecular size of 210 kDa. As indicated in Fig. 1A is the FPLC chromatography of the fractionated TSP sample. Also, as shown in Fig. 1B, the induced lanes showed higher band intensity than the not-induced lanes, indicative of positive induction results.

**Fig. 1.**
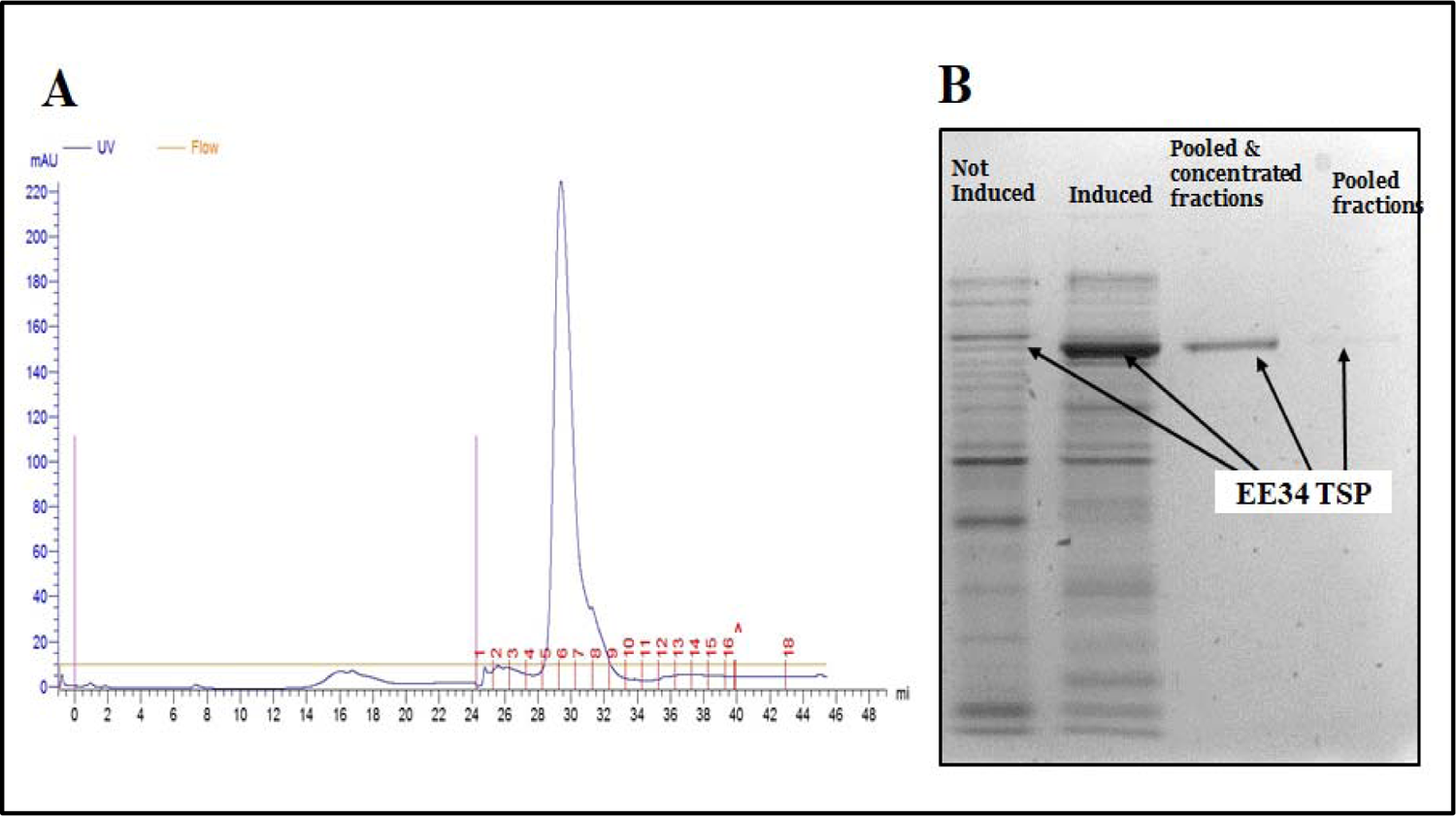
FPLC purification and SDS-PAGE analysis of EE^34^ TSP protein. A) Chromatograph of EE^34^ TSP fractionation. B) SDS-PAGE analysis of fractionated EE34 TSP The 5 mL Cobalt-NTA FPLC columns (Co-NTA) was used in the FPLC fractionation of EE34 TSP samples at pressure of 0.27 MPa, flow rate of 1 mL/minute. Samples that fell under the curve (28-32) were pooled together and concentrated using Amicon Ultra concentrators (Millipore). Samples of both pooled and concentrates were loaded in to SDS gel to analyze for purity. Fractions were eluted with 20 mM phosphate, 400 mM NaCl, 250 mM imidazole solution.

### 3.3. Confirmation of the cloning of the fusion peptide by peptide cleavage

The purpose for the preparation of this fusion tail protein was to add a His-tag protein sequence for protein purification, and once purified, the wildtype TSP (with no His-tag) could be prepared for further study. Furthermore, the peptide could be cleaved off by incubation with recombinant enterokinase (rEK). In this experiment, rEK digestion was utilized to enzymatically cleave the fusion peptide from the purified EE34 TSP, which produced the regular 606-amino acid product from the 649 EE34 TSP; thus, resulting in the production of E34 TSP normal protein with a reduced molecular weight of 196.5kDa instead of 210 kDa uncut protein.

In this study, the presence of the DDDDK protein sequence (the rEK cleavage site) in the putative cloned candidate was demonstrated by incubating the protein with the rEK enzyme for 48 hours and running SDS-PAGE analysis to verify. SDS-PAGE analysis of rEK digestion of EE34 TSP is shown in Fig. 2. Lanes 3 and 4 contained undenatured EE34 TSP, whereas the untreated sample is in lane 3 and the rEK treated sample is run in lane 4. Lane 3, which contains the undigested and unheated samples migrated slightly slower because it contained the 43 amino acid fusion peptides attached to the E34 TSP; however, the third and fourth-lane samples showed a slightly faster shift in migration, this shift is due to the digestion of the 43 amino acid fusion peptides, reducing the molar weight by a factor of approximately 14.1 kDa per trimer. Lanes 7 and 8 display the denatured monomeric species that were treated with rEK (lane 7) and untreated (lane 8). The treated monomeric sample migrated faster than the untreated sample, with consistent loss of the peptide fragment in lane 7. Lysates of undenatured EE34 TSP and denatured monomers from lysates were also studied.

**Fig. 2.**
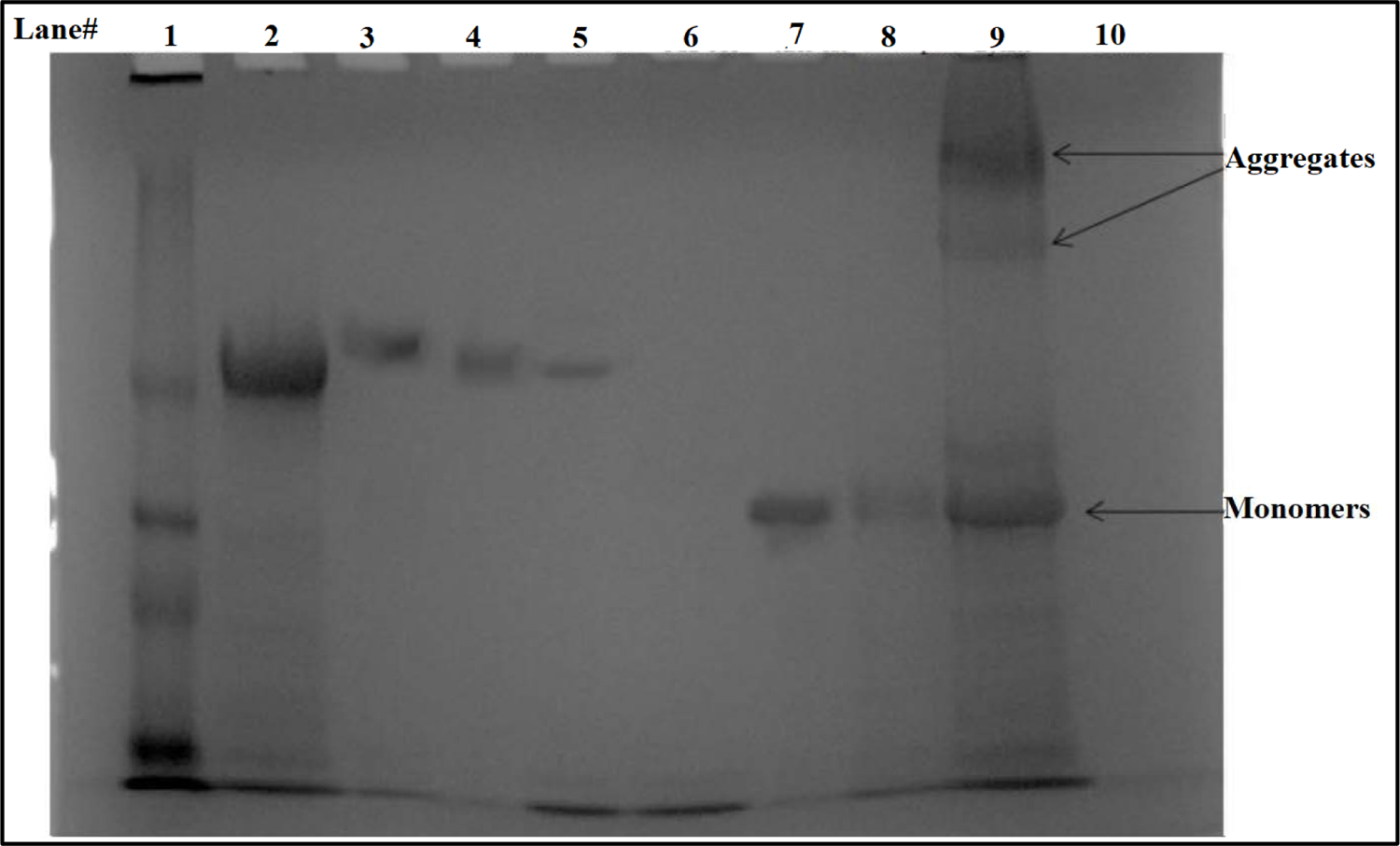
SDS-PAGE analysis of rEK digestion of EE^34^ TSP. Lane 1= Protein standard. Lane 2 = Native EE34 TSP lysate. Lane 3 = Native EE^34^ TSP Pure. Lane 4 = rEK digested pure EE^34^ TSP. Lane 5 = P22 TSP. Lane 7 = Unfolded rEK digested pure EE^34^ TSP. Lane 8 = Unfolded EE^34^ TSP. Lane 9 = Unfold EE^34^ TSP lysate. Lane 10 = Blank

Lane 8 had a slightly slower migration; the difference from samples of the seventh and eighth lanes difference is accounted for by the 4.7 kDa difference in molecular weights between the undigested samples in lane 7 and 8, which were rEK digested to remove the 43-amino acid fusion peptide (43aa is approximately 4.7 kDa).

rEK treated TSP samples are shown to migrated faster than the untreated samples (Figure 2), indicating that the untreated EE34 TSP contained the additional fusion peptide that has region-specific to rEK cleavage site DDDDK. The expressed protein (EE34 TSP) migrated at a size equivalent to 210 kDa protein (196 kDa TSP + 14.06 kDa (three 43aa fusion peptide)). In addition, other studies (data not shown) using a monoclonal antibody to the His-tag sequence (6x-His Epitope Tag monoclonal antibody, Cat. No. MA1-21315-HRP) only identified samples that have not been treated by rEK but could not detect the same samples that had been treated with rEK. This indicated that the EE34 TSPs did consist of the fusion protein, which could later be used for studies necessitating the native E34 TSP.

### 3.4. Differential effect of SDS on P22 and EE^34^ TSPs

Kinetically stable proteins are stabilized in their final state and require large perturbations to change structural features. It has been shown that rigid protein structures containing oligomeric beta-sheets are the basis for kinetic stability, SDS resistance as well as protease resistance of these proteins with such constitutions **(Manning and Colón, 2004; Angela et al., 2018)**. These proteins are known to be trapped by energy barriers **(Manning and Colón, 2004)** in definite conformations in their final native state. P22 TSP, one of the homologous proteins to E34 TSP is highly resistant to SDS unfolding **(Williams et al., 2019)**, and classified as a kinetically stable protein. To investigate for similarity in property between these two proteins, we tested the EE34 TSP against differing concentrations of SDS to verify its resistance or susceptibility to this detergent and to compare with published works on P22 TSP.

The P22 TSP was used here as a control sample since it is resistant to SDS under these conditions. This protein produced a single native-sized discrete band in all concentrations of SDS tested (Fig. 3, lanes 2-5) **(Greenberg et al., 1995; Zayas and Villafane, 2007; Chen & King, 1991; Manning and Colón, 2004)**. In lane 6, in the absence of SDS, EE34 TSP extended tailspike migrates on a native gel in a blurry consistency, but when SDS was added to the samples, even at minimal SDS concentrations as low as 0.05 %, the protein migrates into four distinct, discreet bands labeled B1, B2, B3, and B4. B1 is the slowest moving protein band, and B4 is the fastest moving band.

**Fig. 3.**
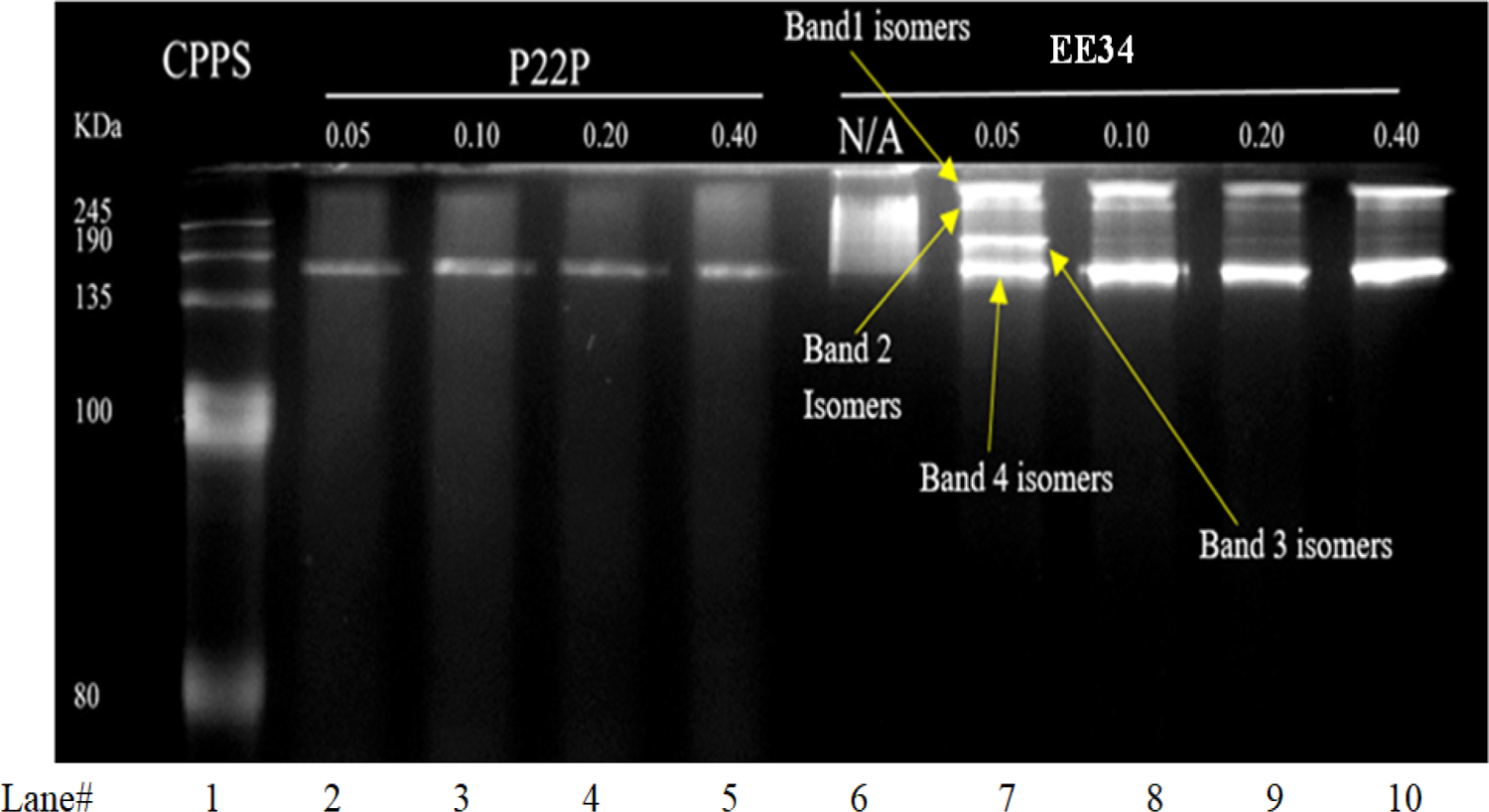
A native electrophoretic gel depicting the migration of extended EE^34^ TSP species under varying concentrations of SDS. Lane 1, Color prestained protein standard; lanes 2-5 are P22 TSP in varying SDS concentrations; lane 6 represents trimeric EE^34^ TSP without treatment to SDS, lane 7 to 10 are trimeric EE^34^ TSP in varying concentrations of SDS.

To quantitatively determine the trajectory of these observed bands, 3 replicate experiments were performed, and densitometric analyses of bands were performed using Image J, and data shown in Fig. 4. It was observed that while the kinetics of each one of these species starts from a common pool of blurry consistency as depicted by a similar densitometric value at 0% SDS concentration. At higher concentrations of SDS, bands 2 and 3 disappeared, and this phenomenon we infer to be that these band species are chased into band 1 and band 4 at the higher SDS concentrations (Figure 3).

**Fig. 4.**
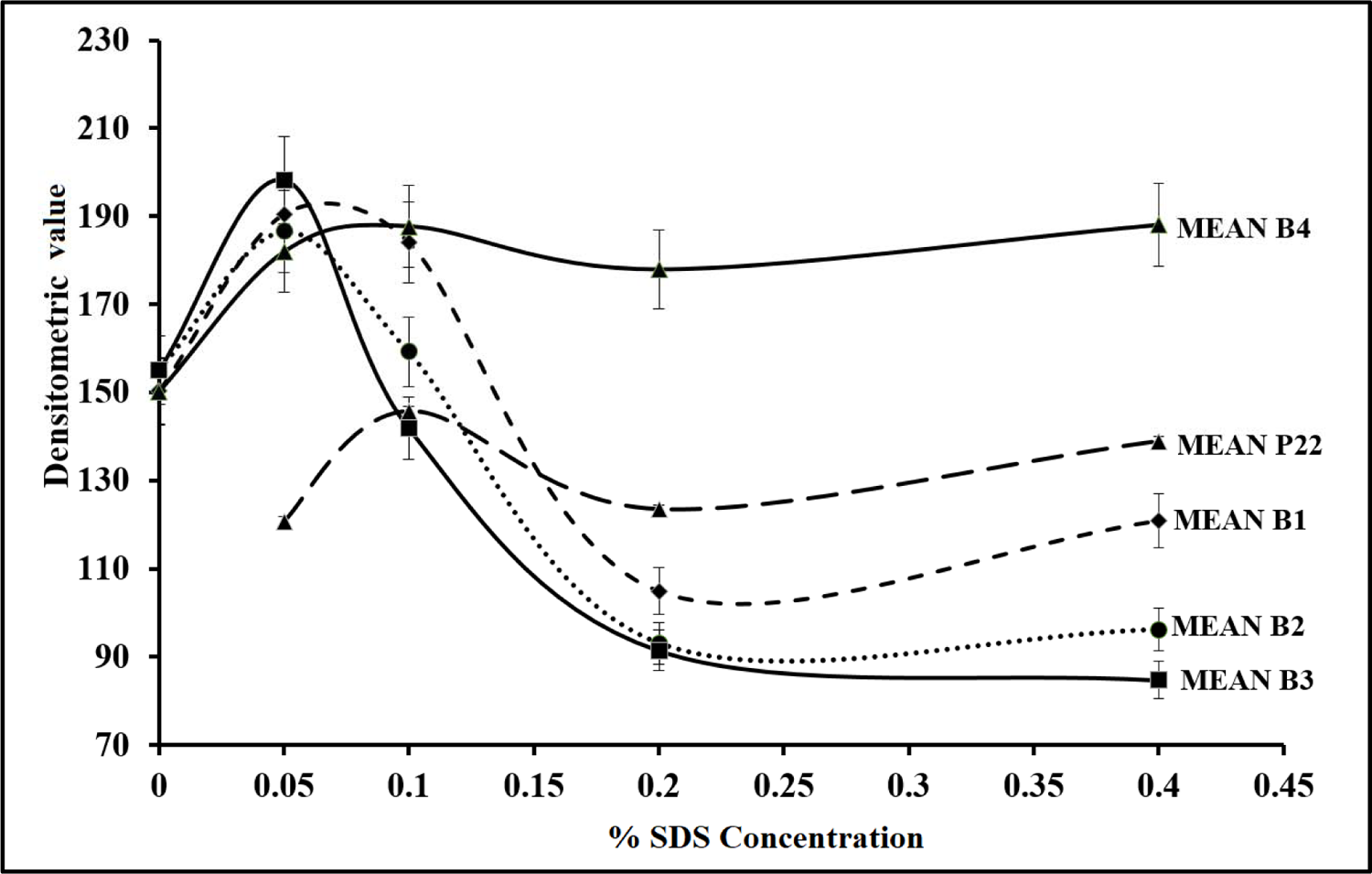
Kinetics of the trimetric isomeric species of EE^34^ TSPs in increasing SDS concentrations. The mean densitometric value of three replicate experiments are presented; all values represent the mean ±SEM (n = 3, p< 0.05). Plots denoted by EE^34^, B1 represents band 1 species; EE34, B2 represents band 2 species; EE^34^, B3 represents band 3 species; EE^34^, B4 denotes band 4 species. P22 tsp was used as a control

In this proposition, the numerous isomers of EE34 TSP were driven into the four main band categories via their interaction with SDS molecules. We propose that B4 protein may be the native conformation of the TSP under these conditions and the most stable. B2 and B3 basically disappear at higher SDS concentrations, therefore less stable. It is to be determined if there is a precursor relationship between these bands.

### 3.5. Thermal denaturation

Incubating protein with heat increases the kinetic energies of the individual atoms that form the molecule. This increase in energy leads to the disruption of bonds used in stabilizing them. Since the stability of the protein in its 3D structure is dependent on weak hydrophobic, hydrogen bonds and electrostatic interactions, increasing heat energy can result in dissociation of these stabilizing bonds **(Ke et al., 2007; Pace et al., 2014; Hatton and Warr, 2015)**. To understand and compare the thermal stability of our proteins which affect such processes such as the virus infection process and its egress from the cell, the EE34 TSP with P22 TSPs were tested by incubation of these proteins to different temperatures of 50 °C, 70 °C, 80 °C and 90 °C alone and then the TSPs were subjected to SDS-heat treatment at similar temperatures. EE34 TSP was incubated at 50 °C under non-denaturing conditions (Fig. 5) and under denaturing conditions (Fig. 6) while Fig. 7 contains the data from incubation under denaturing conditions at three different temperatures (70 °C, 80 °C, 90 °C).

**Fig.5.**
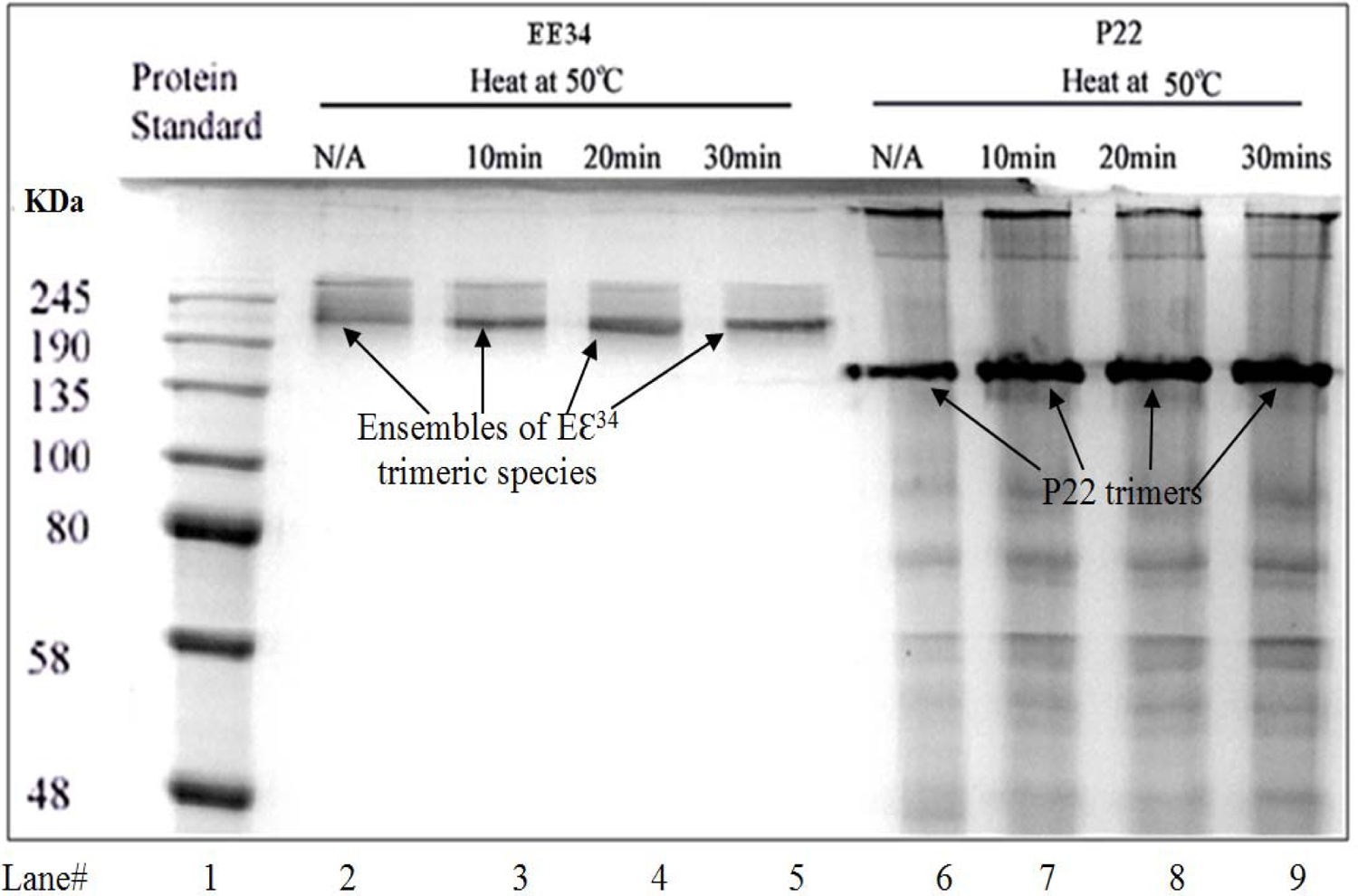
Native gel analysis of EE^34^ TSP and P22 TSP heated at 50 ^°^C. Lane 1, Color prestained protein standard; lane 2 are unheated samples of EE^34^ TSP, lanes 3, 4 & 5 are EE^34^ TSP heated at 50 °C for varying time points; lane 6 is P22 TSP unheated; lanes 7, 8 &9 are P22 TSP heated at 50 °C in varying time points

#### 3.5.1. Incubation of TSPs at 50 **°C** in Native and SDS-PAGE Gels

This study (Fig. 5) displays the effect of 30 mins incubation at 50 °C on the stability of these TSPs under native conditions on EE34 TSP (purified) and on the P22 TSP (from a lysate). This was followed by a similar stability study in which 50 °C incubation was done under denaturing conditions (SDS, Fig. 6).

**Fig.6A.**
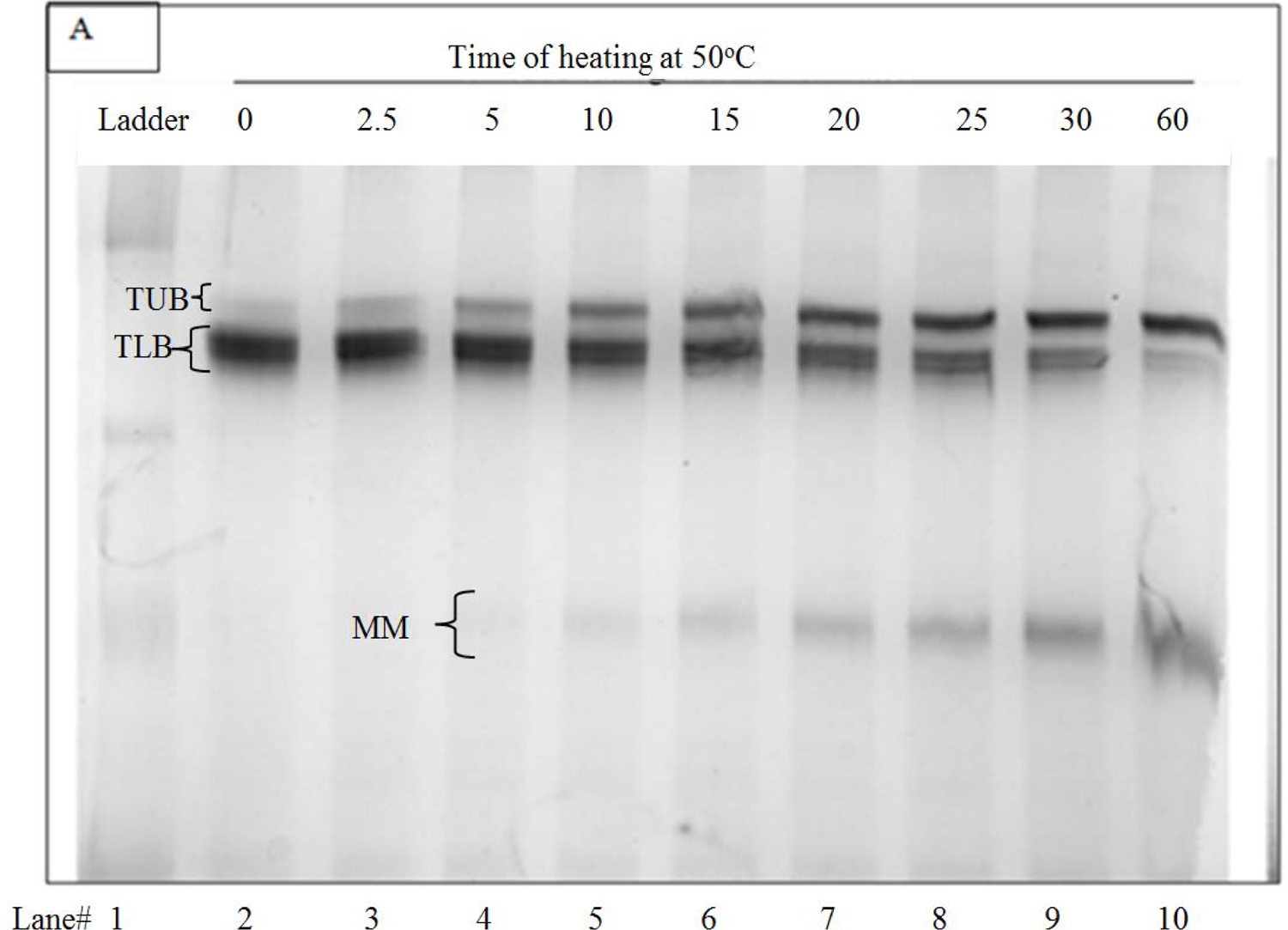
Kinetics of EE^34^ TSP at 50 °C in 0.1 % SDS for an hr. Lane 1, plus stained protein standard; lane 2 are unheated samples of EE^34^ TSP in 0.1% SDS, lanes 3 through 10 are EE^34^ TSP in 0.1 % SDS heated at 50 °C for varying time points

**Fig.6B.**
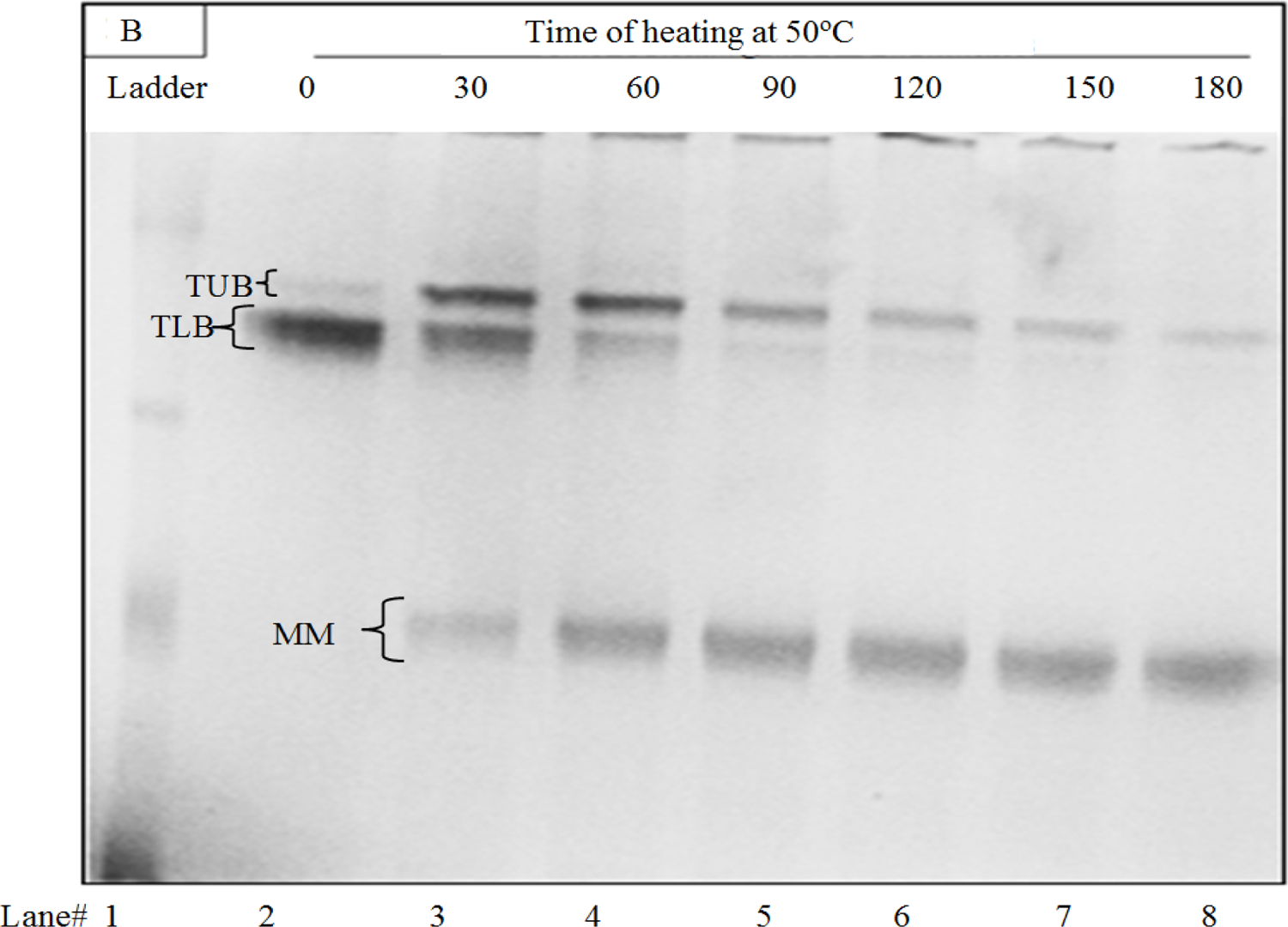
Kinetics of EE^34^ TSP at 50 °C in 0.1 % SDS for 3 h Lane 1, plus stained protein standard; lane 2 are unheated samples of EE^34^ TSP in 0.1 % SDS, lanes 3 through 8 are EE^34^ TSP in 0.1 % SDS heated at 50 °C for varying time points as indicated

Incubating the TSPs at 50 °C (Fig. 5) shows no effect on the EE34 TSP (lanes 2-5) and the P22 TSP (lanes 6-9). Monomers representing the denatured species were not produced by either TSPs at this temperature. The purified EE34 TSP migrated as a multimeric protein ensemble, a “blurry” conformation. The lane 2 labeled “N/A,” which represents the EE34 TSP unheated samples with no SDS, migrated as a blurry conformation in native gels such as observed in Fig. 5 of the SDS study. However, as incubation continues at 50 °C for 10 mins, 20 mins, and 30 mins, it can be seen that there is an accumulation of protein at the bottom of the ensemble as depicted in lanes 3 to 5 of Fig. 5. Also observed, the EE34 TSP migration is noted to be altered since the EE34 TSP is now migrating more slowly than the P22 TSP. Although the molecular weight of the P22 TSP is larger than that of the EE34 TSP, changes in mobility have been observed in the P22 TSP when inserts were placed at its TSP NTD **(Carbonell and Villaverde, 1998)** confirming that the observed change in migration of the EE34 TSP is not an isolated event.

Incubating the P22 TSP at 65 °C in the presence of 0.2 % SDS generated a trimeric intermediate that gets more populated with time. This P22 TSP trimeric intermediate contains a denatured NTD (which consists of the first 108 amino acids of the TSP) and migrates faster than the wild-type P22 TSP. Further incubation results in the formation of a monomeric denatured species from the trimeric intermediate of the P22 TSP **(Chen and King, 1991)**.

In this work, when the EE34 TSP is incubated at 50 °C in 0.2 % SDS for from 60 mins to 180 mins (Fig. 6a and Fig. 6b respectively), three bands were also observed in this study: TUB (trimeric upper band), TLB (trimeric lower band) and MMB (monomeric band). The control EE34 TSP (which was unheated) can be seen in lane 2 of both gels, each of which contained a large thick protein band with a much lighter band above it. We believe the thick TLB represents the native EE34 TSP. The TLB decreased in concentration for the first 60 mins at the same time, the TUB and the monomers (MM) increased in concentration. The largest amounts of these bands (Fig. 6a) occurred at 0 min for TLB, 30 mins for TUB, and at 60 mins for the MM.

To determine the end points at which these bands would be exhausted, the study as in Fig. 6a was performed, and its time was extended for 3 hrs (Fig. 6b). In this study, samples were analyzed every 30 mins for 3 hr. These samples clearly define that TLB decreases as TUB increases. With further incubation, the TUB decreases as the monomer increases. The maximum accumulation of these bands for Fig. 6b occurred at 0 min for TLB, 30 mins for TUB, and after 90 mins for the monomer. These new data on EE34 TSP identify an unfolding mechanism in which the native trimeric species (TLB) are chased to different bands (conformers) represented by a slower migrating upper band species (TUB), which in turn dissociate into monomers with time. Heat treatment seems to convert to a slower migrating species (TUB), a unique finding which contrasts to the mechanism observed in the P22 TSP dissociation **(Chen and King, 1991)**.

#### 3.5.2. Incubation of TSPs at different temperatures under denaturing conditions

To determine the kinetics of the formation of trimers and monomers as well as the appearance of aggregated species near the loading well the study reported as Fig. 7 was performed. Treatment of EE34 TSP, at 70 °C in 0.2 % SDS resulted in the production of a large, diffused band which persisted until 5 mins when only the topmost part of that diffused band remained. By 30 mins, this upper band disappeared, leaving only the monomer migrating around 80 kDa and aggregated proteins near the loading lane, which became intense at 15 mins (Fig. 7a). Treatment of EE34 TSP at 80 °C resulted in aggregation becoming intense at 5 mins, and after that, the monomer became intense at 2.5 mins (Fig. 7b). The 90 °C treatment showed no trimers by 2.5 mins, and aggregation was observed by 2.5 mins (Fig. 7c). Another general trend observed was the decline in monomeric TSP forms, which corresponded to the increase in the aggregation species.

**Fig.7A.**
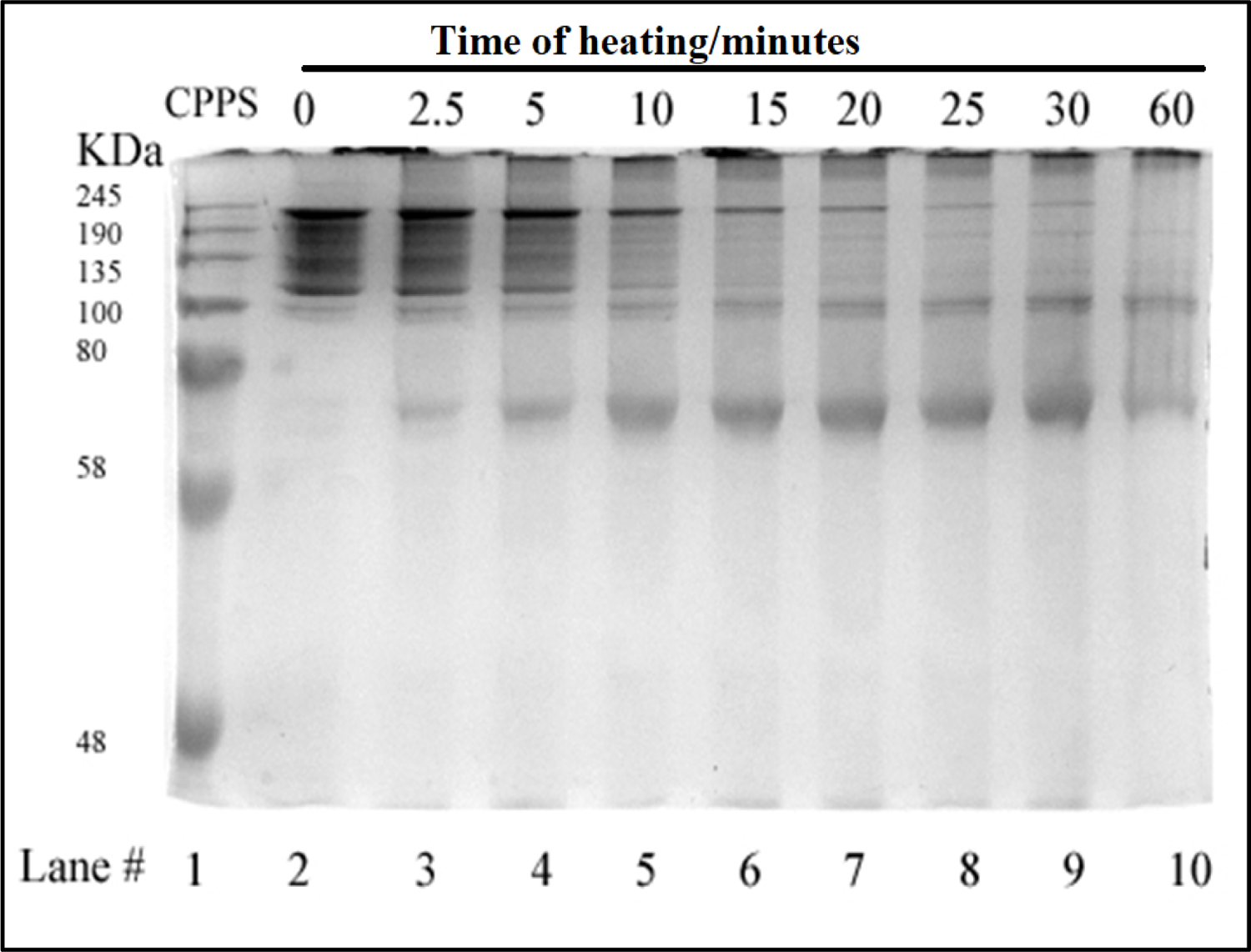
SDS PAGE analysis of EE^34^ TSP at 70 °C.

**Fig.7B.**
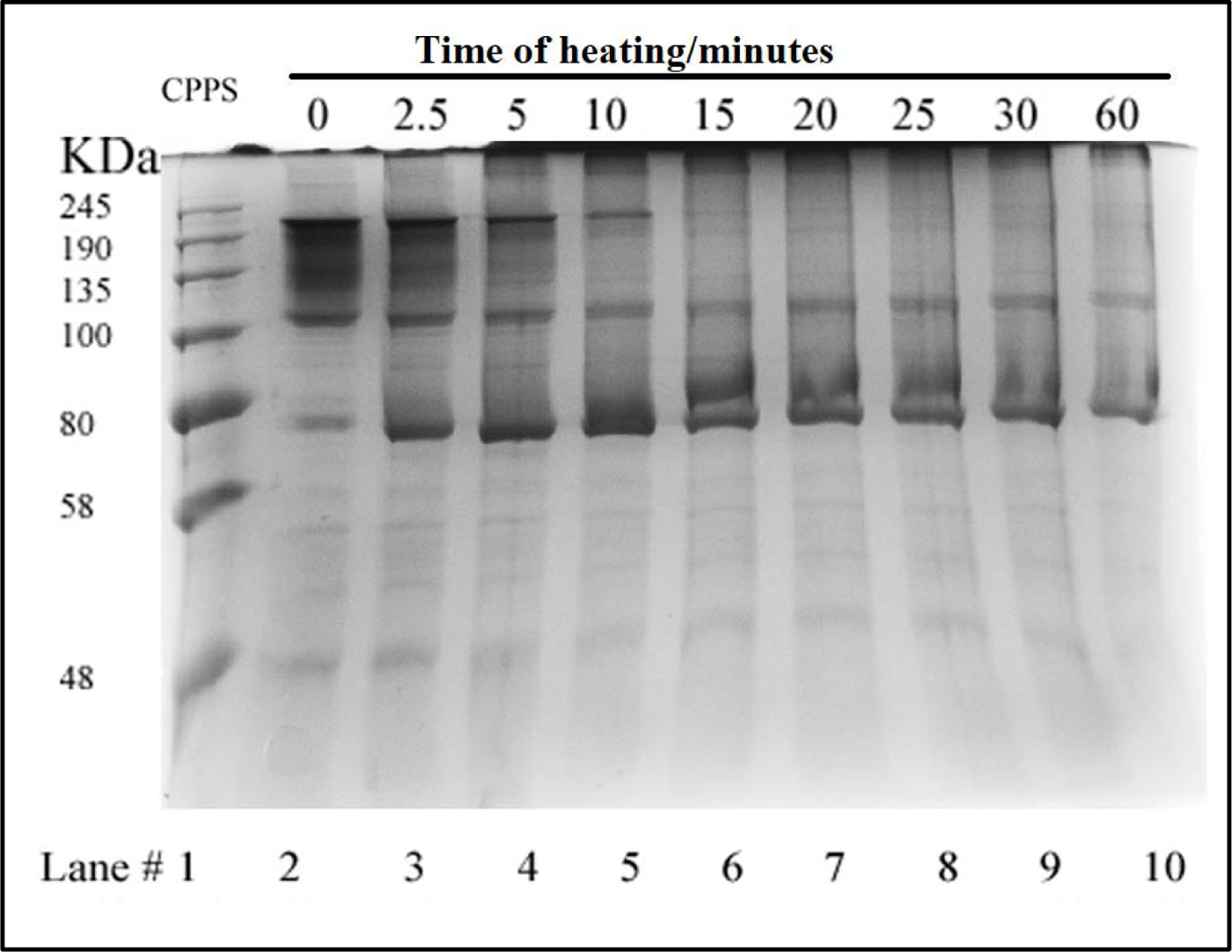
SDS PAGE analysis of EE^34^ TSP at 80 °C.

**Fig.7C.**
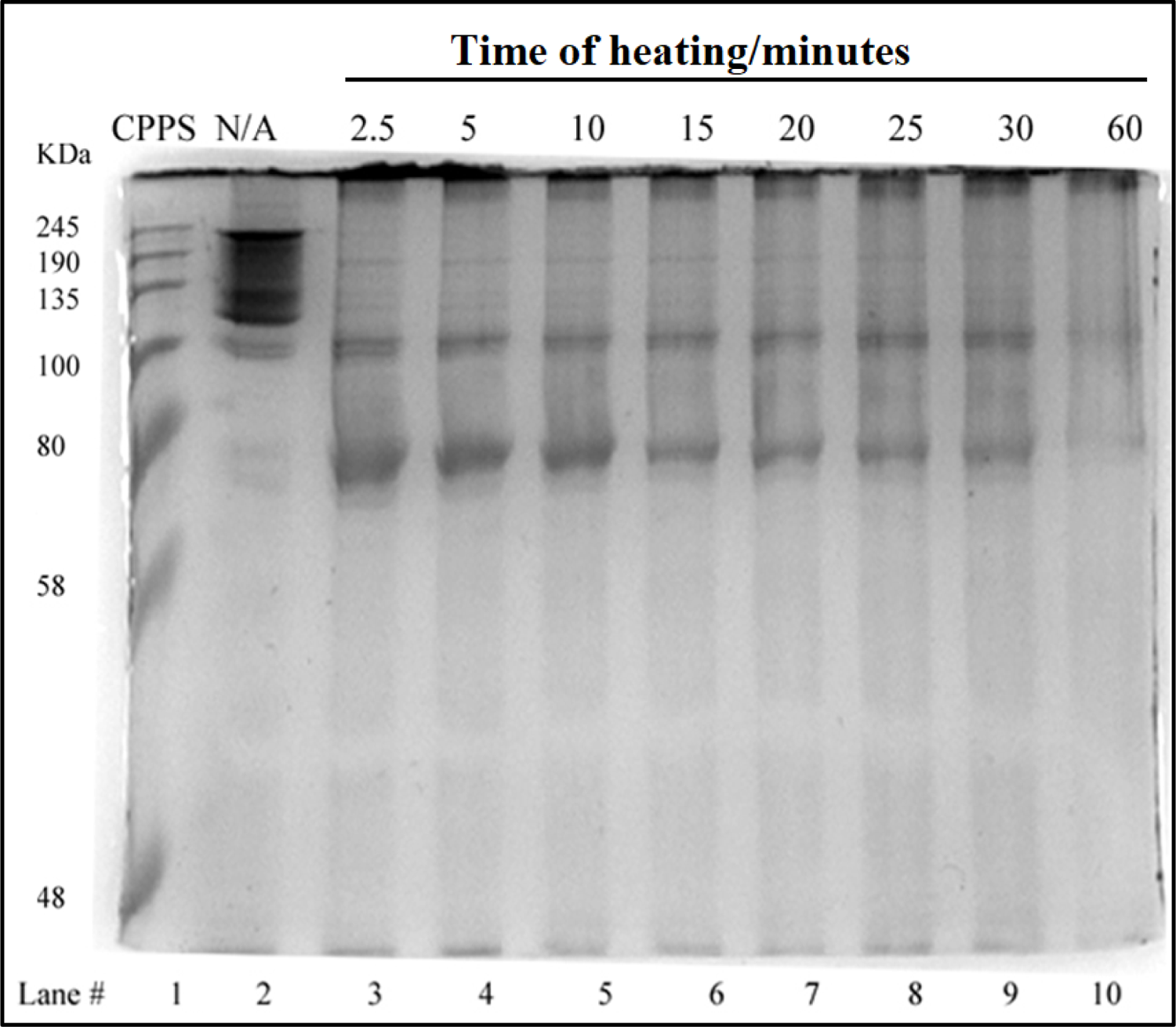
SDS PAGE analysis of EE^34^ TSP at 90 °C. Fig.7A-C. SDS-PAGE analysis of EE^34^ TSP heated at varying temperatures EE^34^ TSP has been characterized at three additional temperatures: 70 °C (Fig. 7a), 80 °C (Fig. 7B), and 90 °C (Fig. 7C) in 0.2 % SDS and analyzed with SDS-PAGE. Under these conditions, the untreated samples form an ensemble of protein bands at all temperatures. The boundaries of these “fuzzy bands” are more intense than the interior of these bands. The internal part of these fuzzy bands disappears upon incubation at these temperatures within 10 min at 70 °C (Fig. 7A, lane 5), within 10 min at 80 °C (Fig. 7B, lane 5) and lastly within 2.5 min 90 °C (Fig. 7C, lane 5). The entire fuzzy band disappears within 30 min at 70 °C (Fig. 7A, lane 9), within 15 min at 80 °C (Fig. 7B, lane 6) and before the earliest measurement at 2.5 min 90 °C (Fig. 7C, lane 3). Denatured EE^34^ TSP (monomers) fastest protein band in every well while bands near the gel top, aggregates are observed after 2.5 min at all temperatures.

#### 3.5.3. Kinetics of monomeric and trimeric species of extended E^34^ TSP in different temperatures

To follow the kinetics of monomers in different temperatures in denaturing conditions, three replicate experiments were performed, and densitometric analysis of bands was undertaken, as depicted in Fig. 8. At the zero-time point, all monomers showed a very low abundance at all temperatures studied, each registering less than 20 %. After 2.5 minutes of heating at 70 °C, 80°C, and 90 °C, it produced drastic changes in monomeric species abundance. While a drastic change is recorded with the 90 °C at 2.5 minutes, resulting in a 97.9 % monomeric E34 TSP population, 70 °C seemed to produce no significant effect on the protein at this time point. Heating at 80 °C produced 57.1% monomers.

**Fig.8A.**
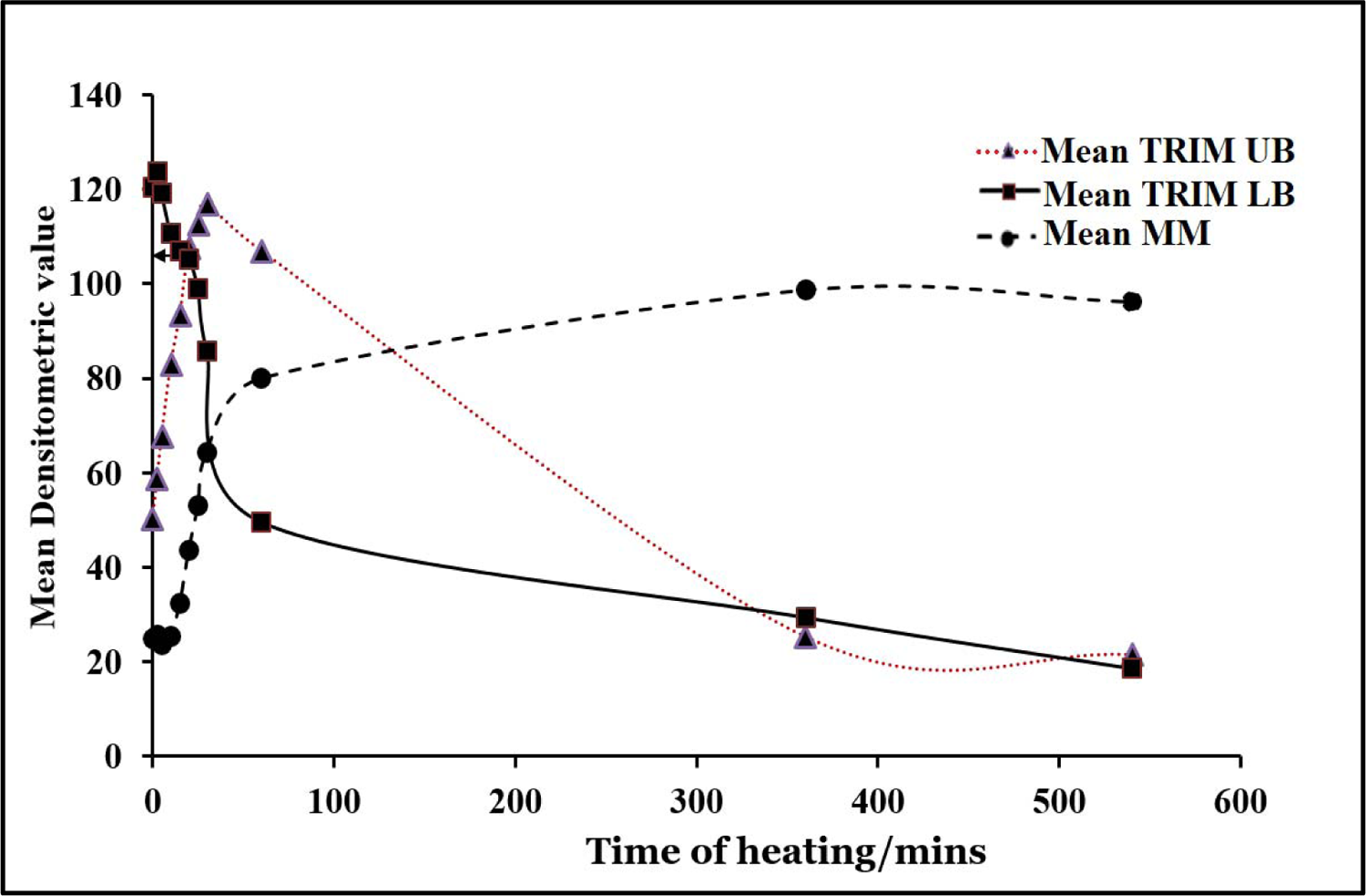
Thermal unfolding kinetics of EE^34^ TSP at 50 °C.

**Fig.8B.**
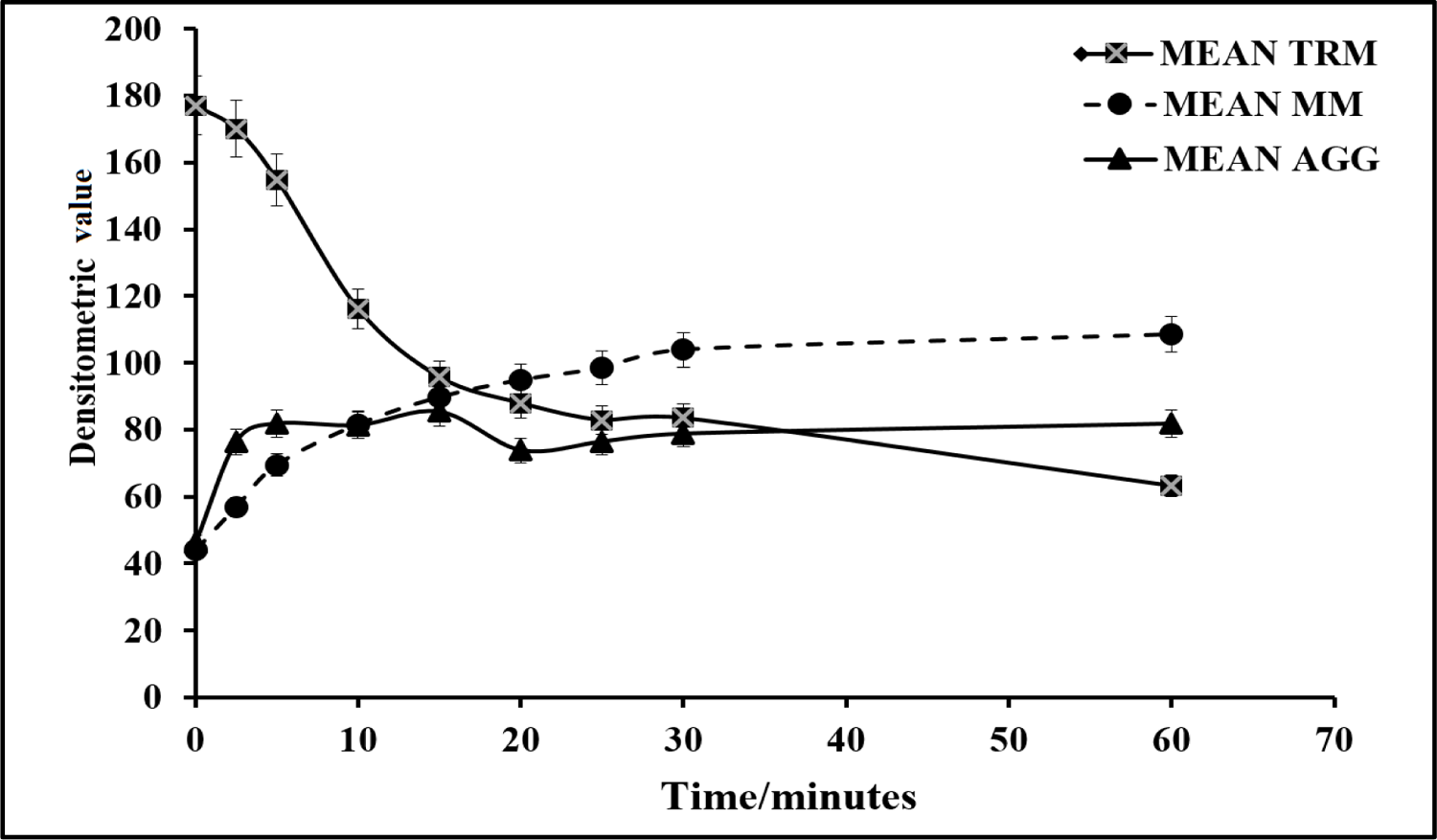
Thermal unfolding kinetics of EE^34^ TSP at 70 °C

**Fig.8C.**
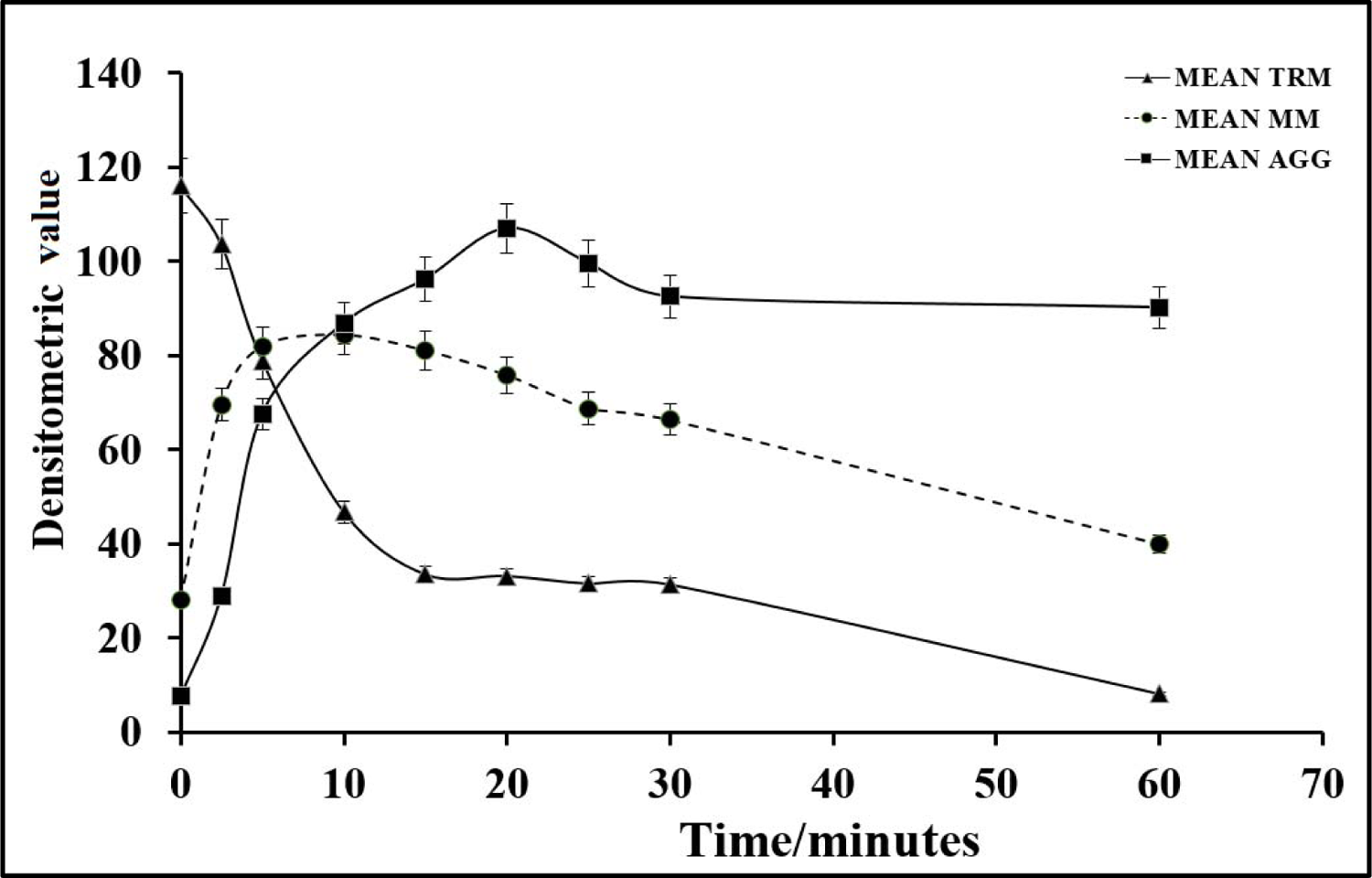
Thermal unfolding kinetics of EE^34^ TSP at 80 °C

**Fig.8D.**
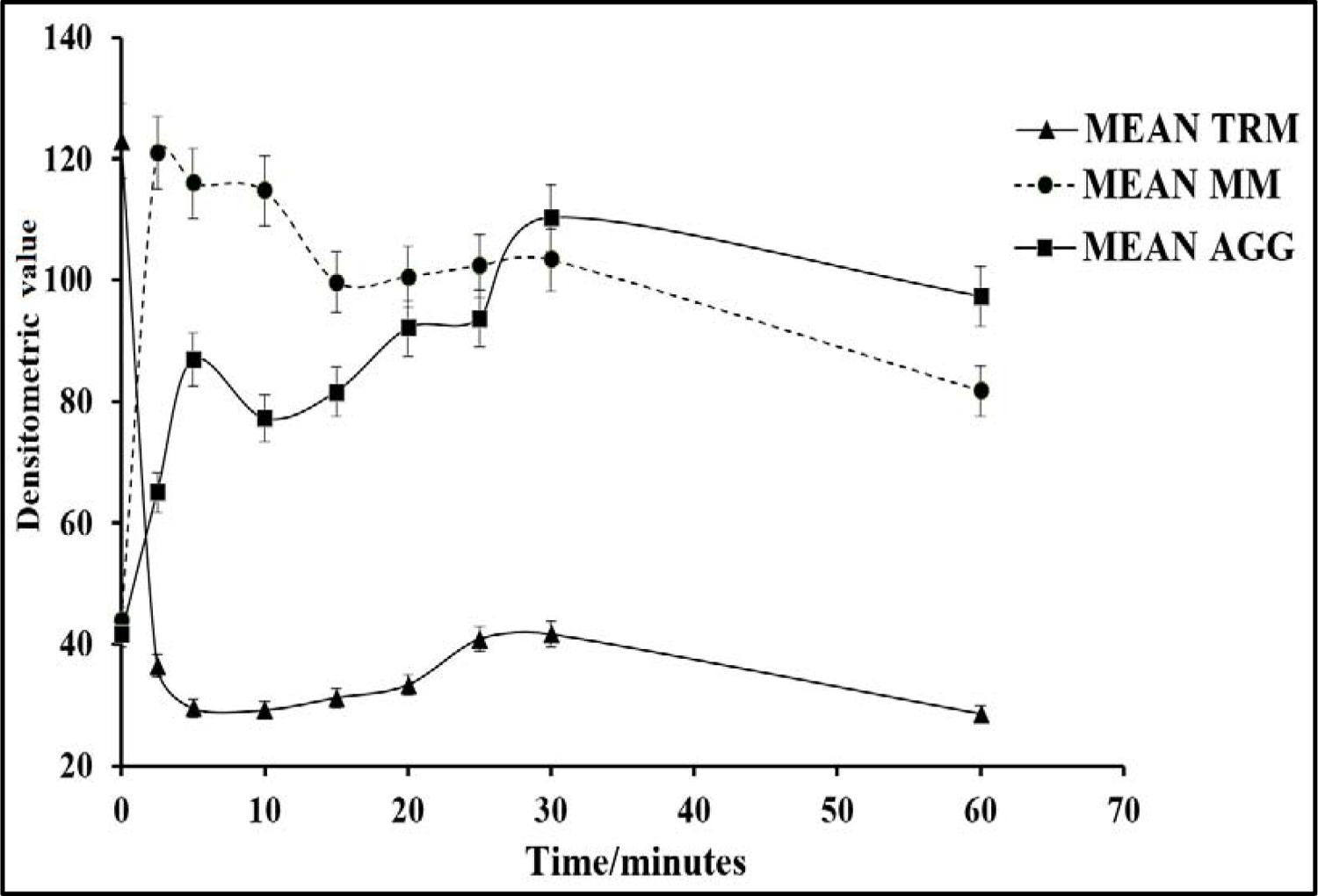
Thermal unfolding kinetics of EE^34^ TSP at 90 °C Fig. 8A-D. Thermal unfolding kinetics for EE34 TSP under different temperatures (50 °C, 70 °C, 80 °C and 90 °C), mean TRM; mean trimers, Mean MM = Mean monomers, Mean AGG = Mean aggregates. Mean TRIM UB = Mean trimers upper band species of EE34 TSP, Mean TRIM LB = Mean trimers Lower band species of EE34 TSP. Graphs are plotted as mean values and standard error of the mean (bars) from three independent experiments.

As shown in Fig. 9, after 5 mins, heating at 80 °C and 90 °C produced over 65 % and 90% monomers, respectively. The highest percentage of monomers produced at 80 °C is observed at the 10-min mark, while the highest monomeric species by heating at 90 °C came just 2.5 mins. At 70 °C, it took an hour to produce its highest abundance of monomers, at just 47.4 % abundance. Meaning, even after an hour of heating at 70 °C, the heat energy generated was not enough to denature the extended E34 TSP above 50 %.

**Fig. 9.**
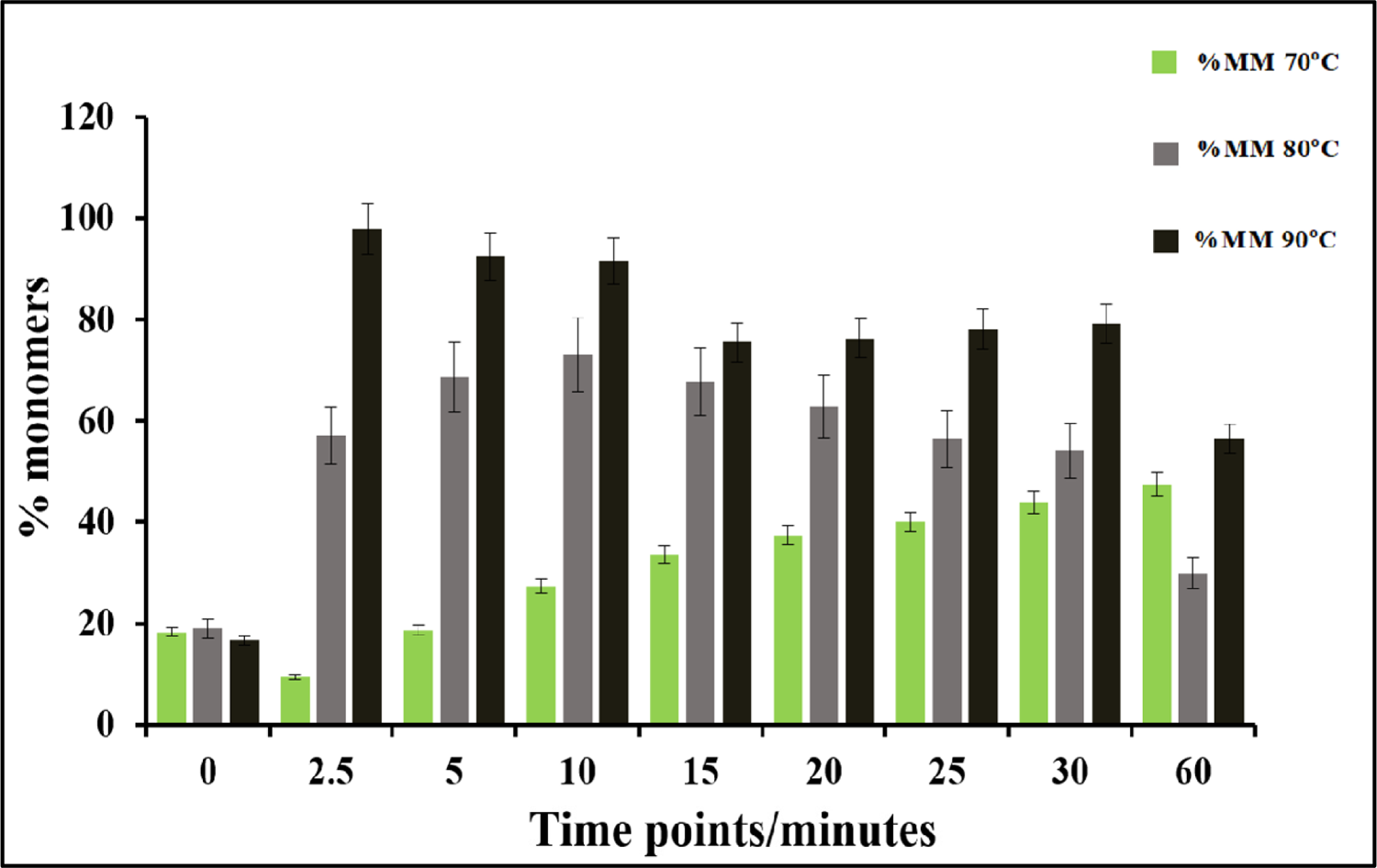
Bar plot of EE^34^ TSP monomers at different temperatures. This chart compares the monomeric species from three different treatments; heating of EE34 TSP in 0.1 % SDS at 70 °C, 80 °C and 90 °C. Green bars represent 70 °C, grey bars represent 80 °C and black bars represent 90 °C. All data presented are derivative of triplicate experiments; all values represent the mean ±SEM (p < 0.05).

One major observation in both the 80 °C and 90 °C experiments was the reduction in monomeric species after the peak abundances were reached. For instance, at 2.5 mins, at 90 °C produced a staggering 97.9 % monomeric species, then, the abundance of monomers gradually and continually decreased with time until a lowest of 56.5% was recorded at 1 hr. While at 80°C, after recording the highest percentage abundance of 73.1% at 10 mins, there is a gradual and consistent decrease in monomeric species percentage until it stood at a minimum of 29.9% at 1 hr.

The percentage abundance of the trimers (Fig. 10) at the start of the experiments for all temperatures stood close to 100 %, the first drastic change in trimer abundance occurred at 2.5 mins for the 90 °C, reducing to a low of 8.9 % and then continued to decrease until it hits the lowest at the 60th minute with an abundance of 0.5 %. It showed the steepest decline in all the heat curves generated. The 80 °C treatment produced a sustained constant reduction and finally recorded 0.46 % abundance in an hour.

**Fig. 10.**
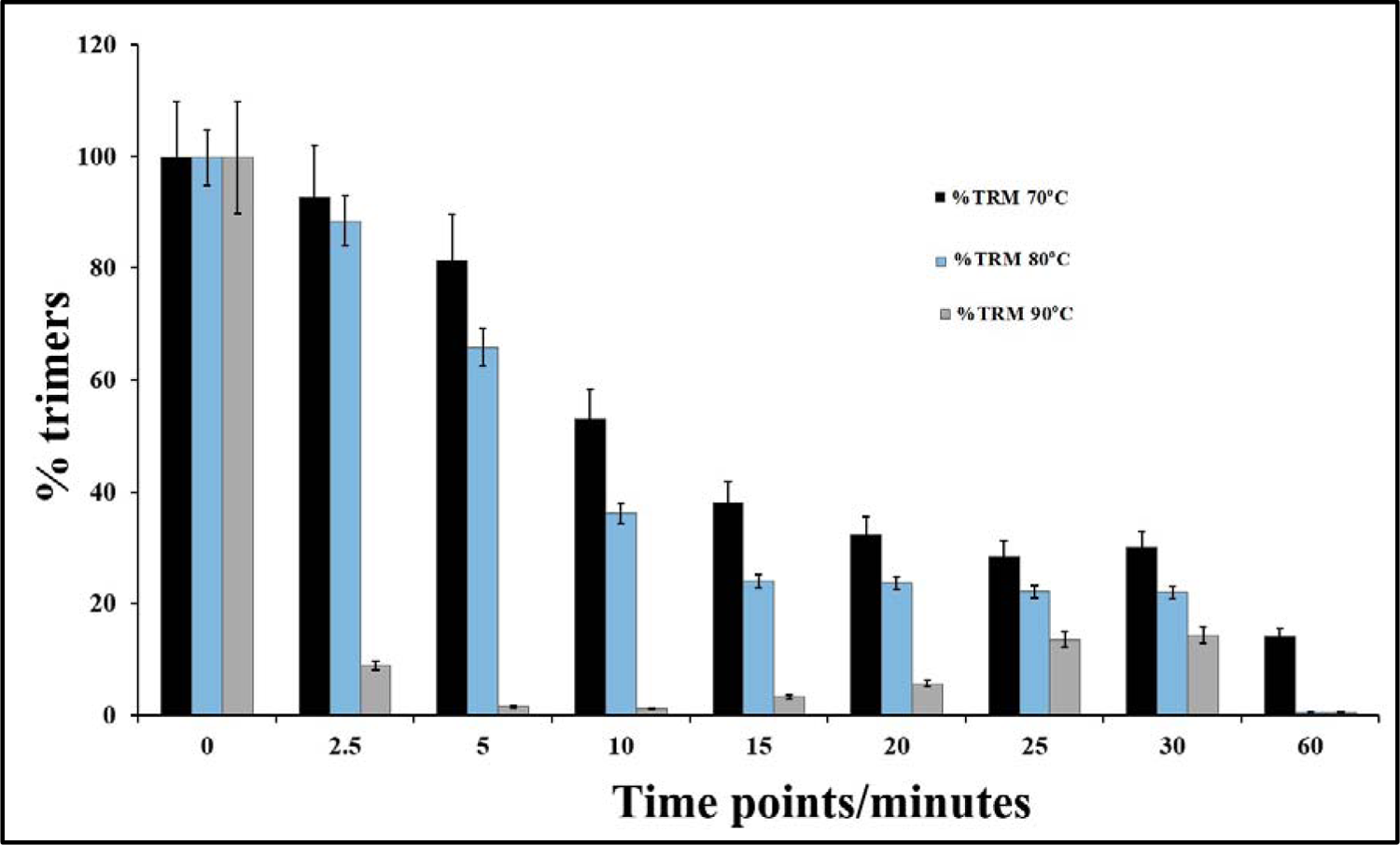
Bar plot of EE^34^ TSP trimers at different temperatures This chart compare EE34 TSP trimeric species from three different treatments; heating of EE34 TSP in 0.1 % SDS at 70 °C, 80 °C and 90 °C. Black bars represent 70 °C, blue bars represent 80 °C and grey bars represent 90 °C. All data presented are derivative of triplicate experiments; all values represent the mean ±SEM (p < 0.05).

Like the 80 °C treatment, the 70 °C treatment produced a similar curve except in comparison; it produced a less steep slope. This gentle slope led to the lowest percentage abundance at the 60 mins of 14.0 % trimers at 60 mins, which indicates an over 80 % decrease in trimeric forms.

#### 3.5.4. Half-life of EE^34^ TSP in different temperatures in the presence of 0.1 % SDS

In measuring the time at which the unfolded and native states of the TSPs have populated equally in solution at each set temperature, we found that the longest time was recorded for the 50 °C heat at 125 mins, and the shortest 0.48 min recorded for the 90 °C indicating that this protein is sensitive to elevated temperatures.

### 3.6. Test for activity

As demonstrated by Fig. 11, the formation of infectious P22 phage is a multistep process involving specific protein-protein interactions. The last component in the assembly pathway is the addition of the highly specialized bifunctional hydrolase (the P22 TSP) for adsorbing the phage to the LPS to the membrane and subsequent infection of *Salmonella typhimurium* **(Ackermann, 1998)**. Since the P22 TSP shares a similar structural topology with E34 TSP, and at the N-terminus, 70% identity with E34 TSP **(Villafane et al., 2008)**, this implies that our EE34 TSP if it had retained the same structure in the N-terminus should also bind to P22 H. As depicted in Fig. 12A, P22 phage heads alone cannot infect its host. When P22 H bind to EE34 TSP, the prosthetic phage cannot infect the P22 phage host (Fig. 12B). Finally, only P22 H bound to P22 TSP are infective hence will bind to P22 host cells, hydrolyze the LPS and inject its DNA into the host (Fig. 12C, D). This understanding was therefore utilized in testing the activity of the EE34 TSP as described below.

**Fig. 11.**
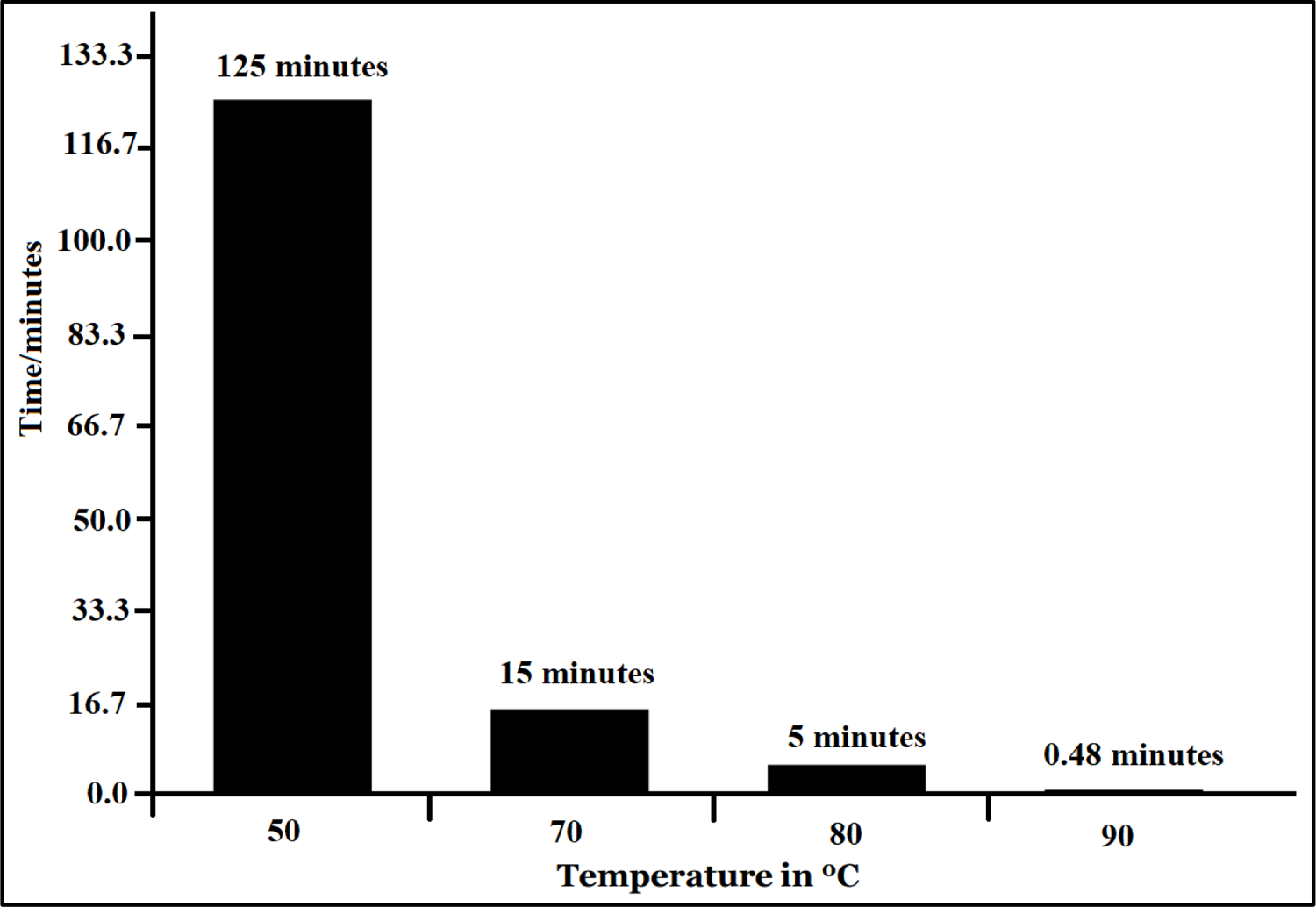
Bar plot of unfolding midpoints of EE^34^ TSP in various temperatures All data presented are derivative of triplicate experiments base on the calculations of the midpoints of the unfolding curves in the graphs generated in Fig. 8A-D. The slowest to reach the unfolding midpoint was EE34 TSP in 0.1 % SDS heated at 50 °C, the faster was 90 °C.

**Fig. 11.**
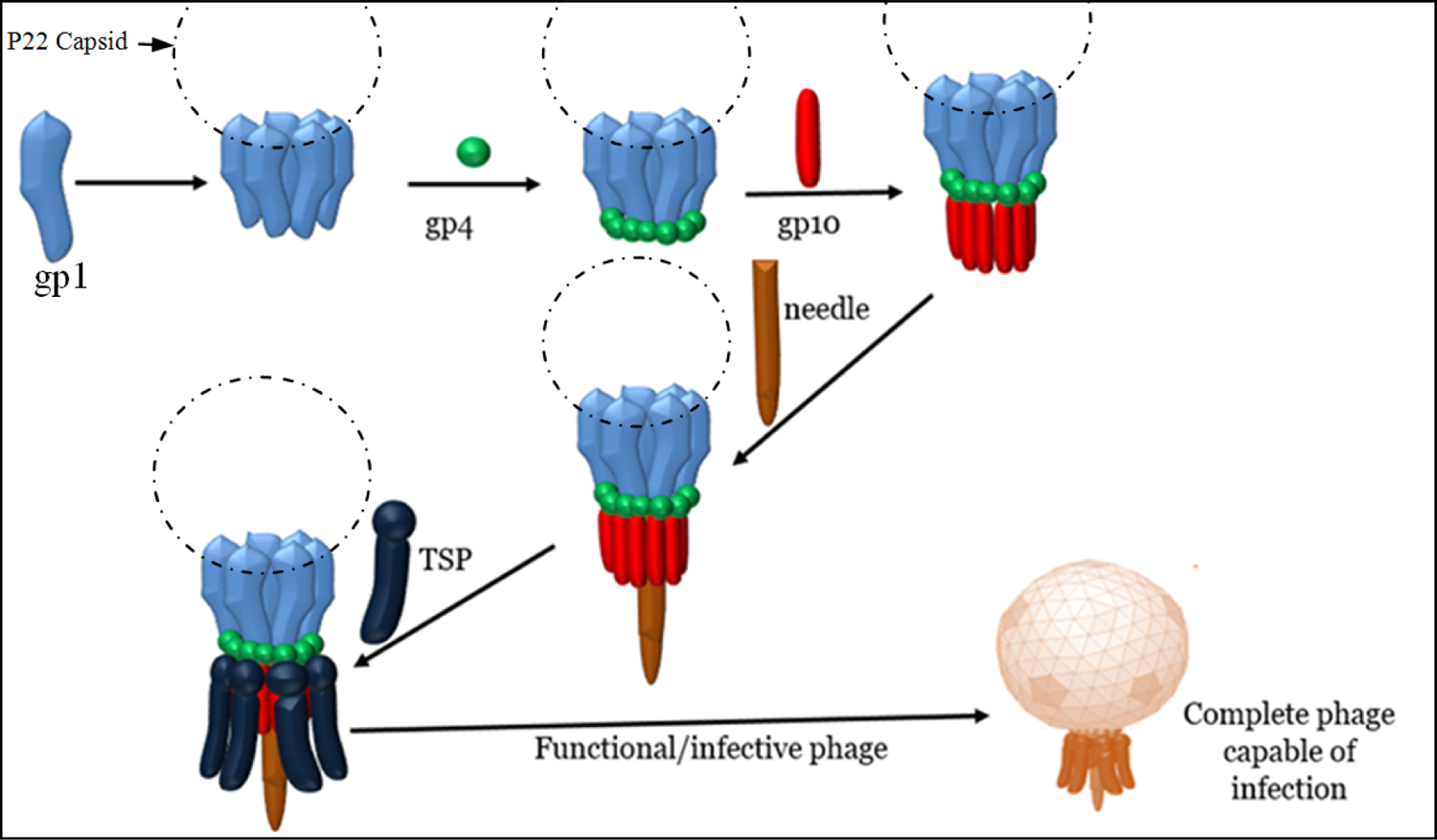
Schematic cartooned diagram illustrating the assembly mechanism of the tail machinery of the P22 phage The P22 tail machine, a fifty-one subunits complex arising from five different gene products is the last assemble process in the maturation of the P22 phage. The portal complex that fit into the capsid of the phage is shown as blue, and following DNA packaging, the tail machine assembles *through a series of* accumulation of many copies of gp4, gp10, the needle protein and the tailspike protein to the portal ring (as indicated by the blue color), the very last step however is the addition of the LPS binding and hydrolase moiety, which is the TSP of the phage. The addition of the LPS binding and hydrolase moiety activates the phage’s infectivity.

**Fig. 12.**
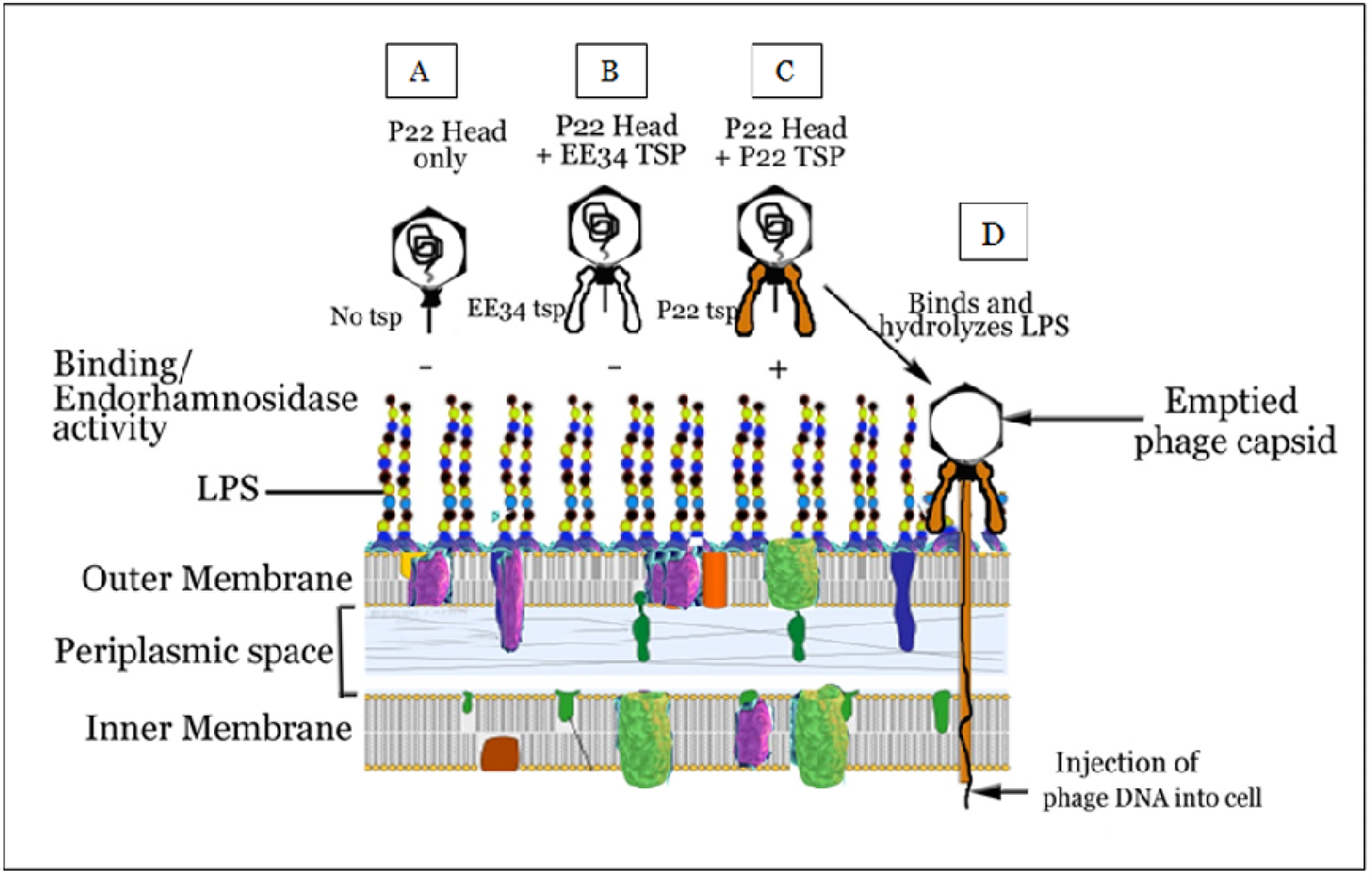
Schematic illustration of the interference assay A= Tailless P22 phage (also termed as the P22 phage heads) which consist of all the phage’ components assembled together except the tailspike protein, B= Prosthetic phage consisting of P22 phage head bound to EE^34^ TSP. C = P22 phage heads bound to P22 TSP to form a complete phage which can bind to the LPS. D = the infective P22 phage which has endorhamnosidase activity on the LPS of the S. typhimurium host leading to DNA injection into the cell and subsequent infection.

#### 3.6.1. Interference of EE^34^ TSP in P22 Phage assembly

In Fig. 13, the result shows that the P22 H - P22 TSP lane (positive control) had spots showing an area of lysis, (i.e., the first three spots with a higher concentration of P22 TSP. Four spots were visible in the P22H-EP22 TSP lane, indicating that the 43-amino cloned variant of the P22 TSP did not affect the protein’s binding to the P22 H to form infective phages. The P22H-EE34 TSP lane showed no area of lysis, indicative of prosthetic phages that could not infect the P22 host. Finally, the P22H-EE34 TSP, P22 TSP lane showed plagues at the last two spots at the right. As indicated by the arrow, concentrations of EE34 TSP at those two spots were low, hence they did not bind to all available P22 H. The residual P22 H could then interact with P22 TSP that was subsequently added after 30 mins.

**Fig. 13.**
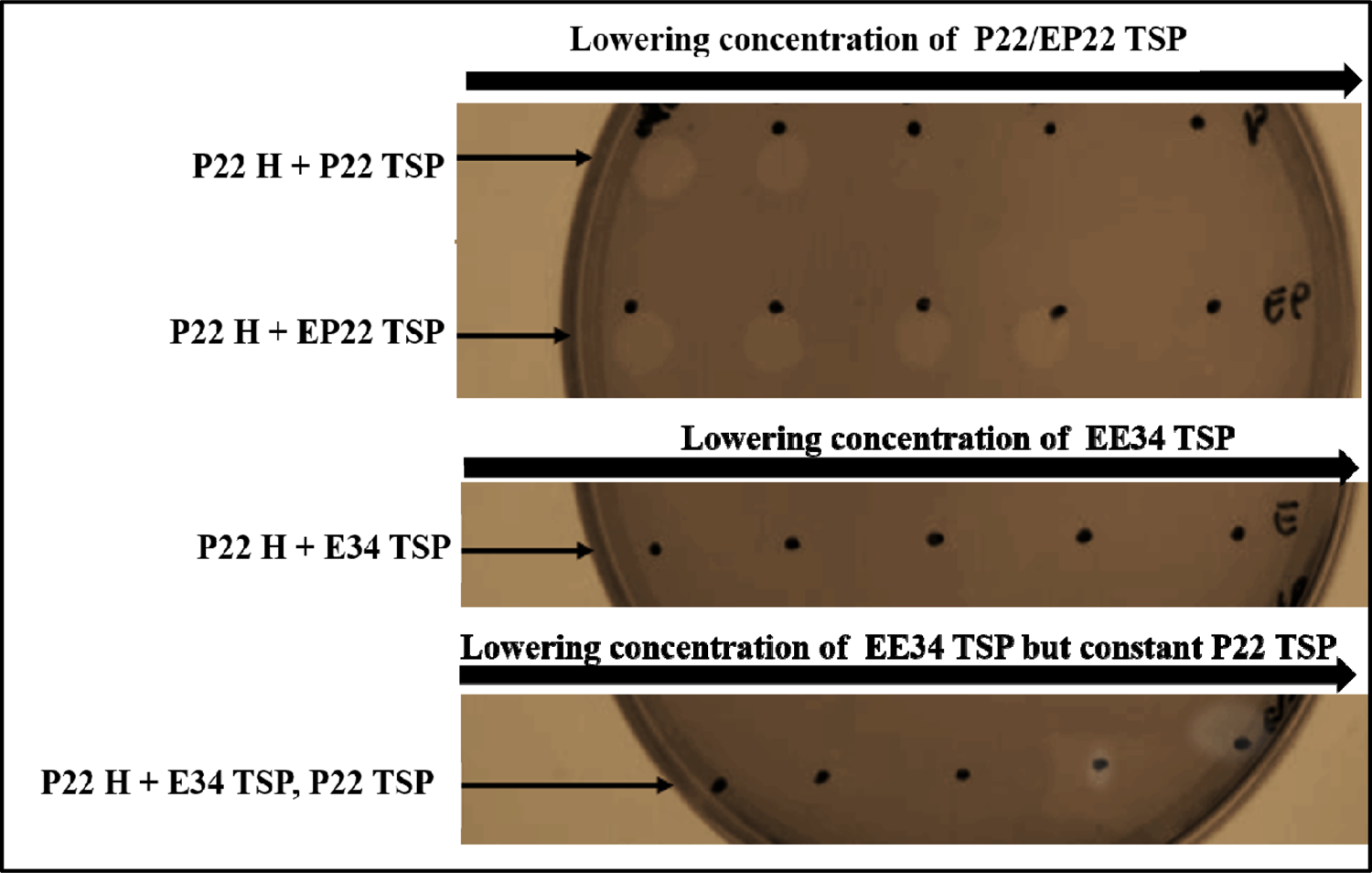
Interference of EE34 TSP in *in vitro* P22 Phage assembly and thereby blocks infectivity. Same concentrations of P22 heads (0.078 mg/mL) were first incubated with EE34 TSP (0.48 mg/mL) for 30 mins, followed by varying concentrations of P22 TSP (2.0, 1.0, 0.5, 0.25, and 0.125 mg/mL) was added to the designated treatment mix and allowed to sit under room temperature for another 30 mins. The P22 TSP-P22 Head (H) row which serves as a positive control received buffer instead of EE^34^ TSP, then followed by the addition of the P22 TSP. Afterward drops of the reaction sample were taken from each treatment and spotted on a lawn of *S. typhimurium* grown on agar plate. The plates were incubated in 37 °C overnight and then photographed.

#### 3.6.2. Quantitative analysis of interference of EE^34^ TSP in P22 Phage assembly

As shown in Fig.14, the effectiveness of EE34 TSP to interfere in the P22 phage heads and P22 phage TSP interaction and hence binding competitively is clearly demonstrated. The negative control represented P22 H treated to buffer only, while the positive control represents P22 H treated with buffer and counter titrated with P22 TSP. The maximum numbers of plagues were recorded for the positive control (and an average of 136.6 plagues from 10 replicates), while almost all negative control plates registered no plague (except a misnomer which came as eight plagues in a single plate of the ten replicates). At the higher concentrations of EE34 TSP, very low counts of plagues were recorded. At the lower concentration of EE34 TSP, an increasing number of plagues were recorded. The stock EE34 TSP sample with a 20 mg/mL concentration to the 5th dilution with a concentration of 0.0002 mg/mL recorded plagues from 1.6 to 17. The 6th serial dilution (0.00002 mg/mL) recorded a high of 33.6 plagues.

**Fig. 14.**
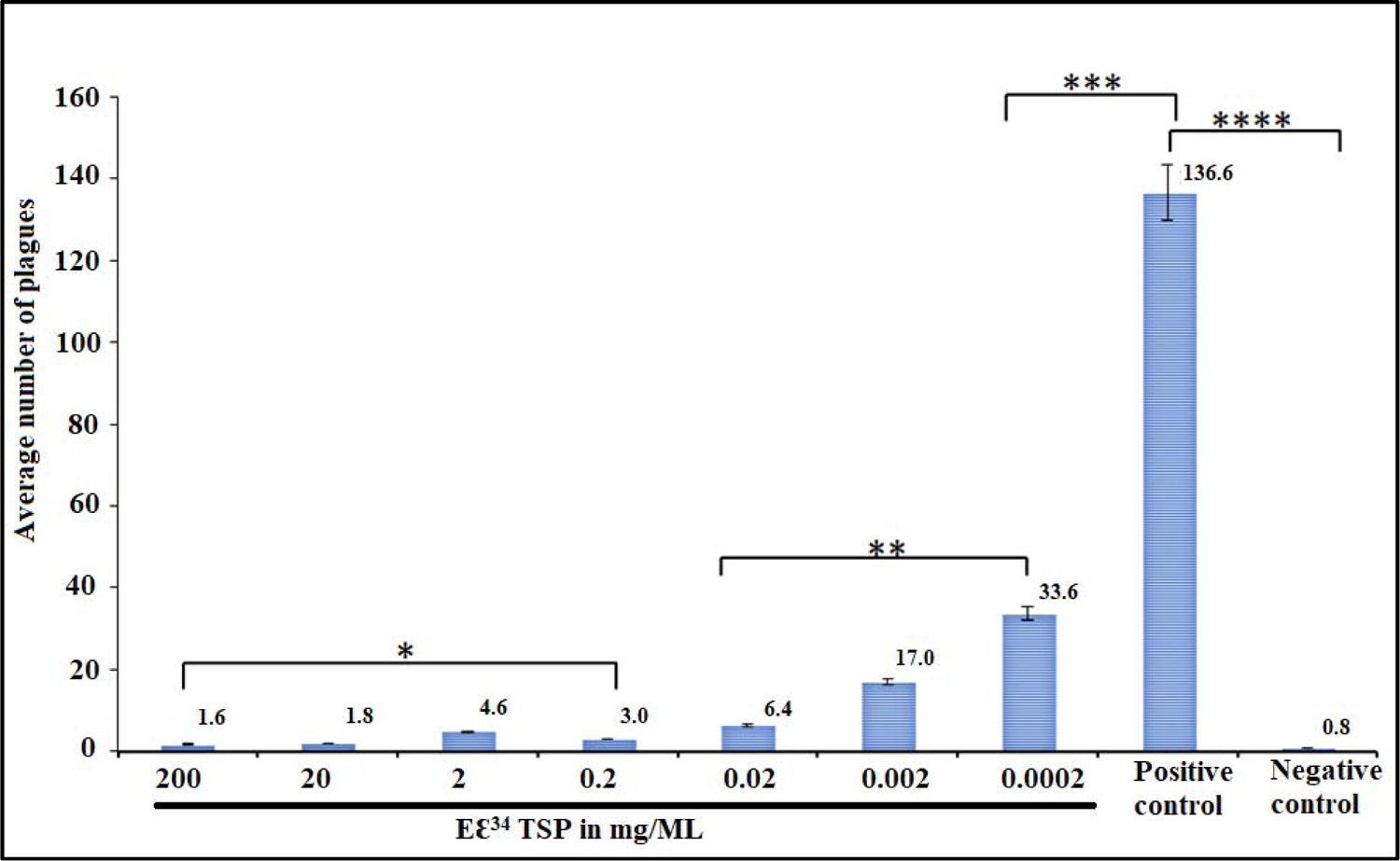
Chart showing the effectiveness of EE34 tsp to competitively interfere in the P22 phage heads and P22 TSP protein interaction. The EE34 tsp ability to competitively interfere in the P22 phage heads-P22 tsp interaction was analyzed via interference assay. The data represents mean ±SD of ten independent experiments. The asterisk indicates statistical significance (* = P>0.2, ** = P>0.1, *** = P>0.05, ****= P>0.0005)

#### 3.6.3. Homology modeling of E34 TSP

The wildtype E34 TSP amino acid sequence from the Gene ID: 7353089 was extracted from the NCBI website^7^. Homology modeling was carried out following a similar process published by Filiz et al., 2014 **(Filiz and Koc, 2014)**. In brief, the sequence was blasted against PDB to extract the most suitable templates for homology modeling. The bifunctional P22 TSP, a homotrimer with PDB ID 2XC1.1.A **(Seul et al., 2014)** possessing 69.83% amino acid identities, was used to model the head binding domain (HBD) of the E34 TSP, whereas the exopolysaccharide biosynthesis protein (bacteriophage tailspike-like parallel beta-helix fold, a homotrimer) of Pantoea stewartii with PDB ID 6TGF.1.A **(Irmscher et al., 2021)** that shows 29.29% identity was used for the homology modeling of the receptor-binding domain of the E34 TSP. As shown in Figure 15A, the HBD showed a globular structure, with the linker region exhibiting alpha-helix turns. This is similar to the crystal structure of the P22 TSP head binding domain as reported by Steinbacher et al., 1997 which published that the P22 TSP possesses a trimeric dome-like structure created by two perpendicular beta-sheets of five and three strands (Steinbacher et al., 1997).

**Figure 15.**
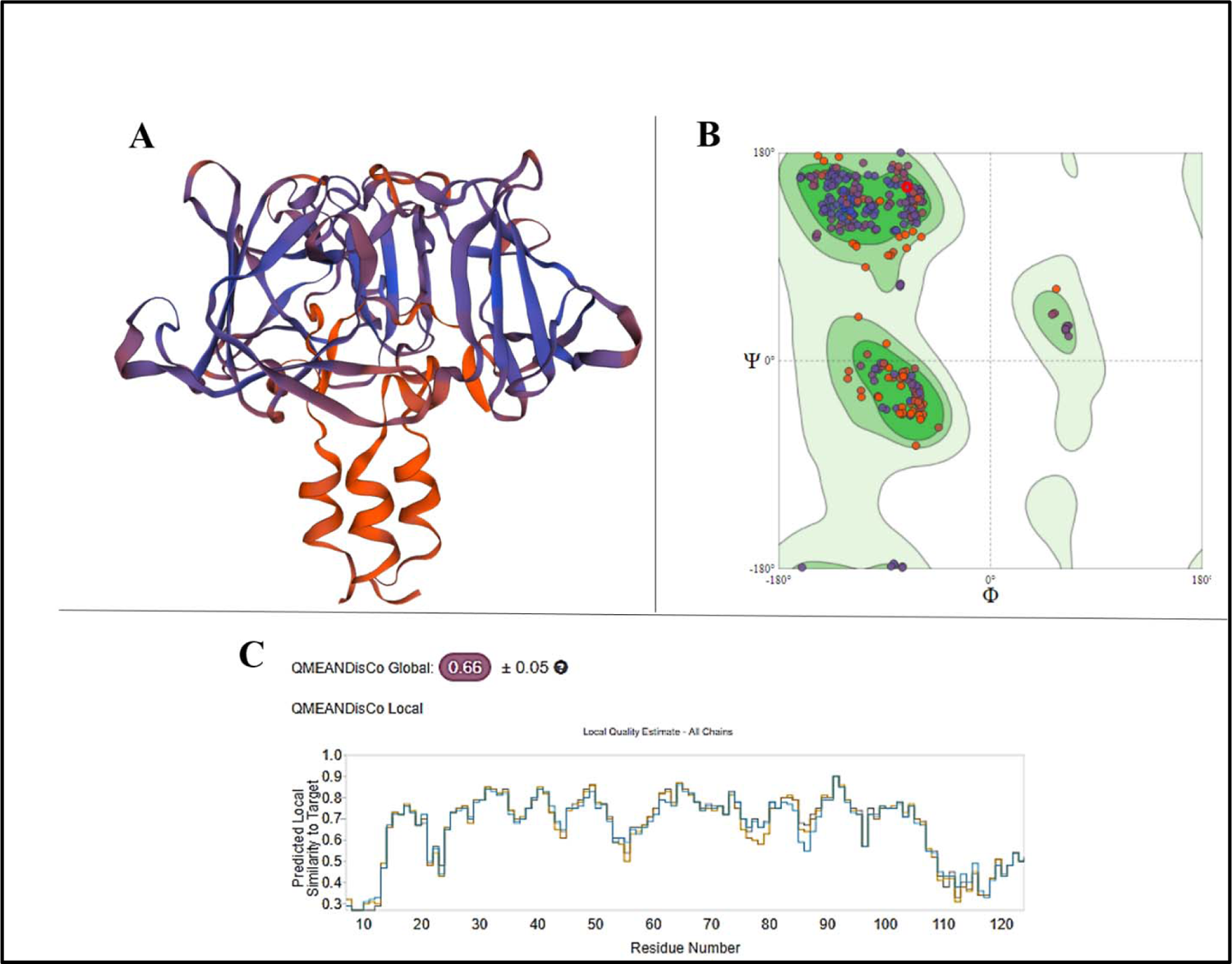
Homology model of the Head Binding Domain of E34 TSP. Using the Swiss-Modeler, the head binding domain (HBD) of the E34 TSP (Figure 15A), which consist of the firs 109 amino acids at the N-terminus were modeled and its stereo-chemical stability evaluated using Ramachandran plot (Figure 15B). PDB ID 2XC1.1.A (Seul et al., 2014) possessing 69.83% amino acid identities, was used to model the HBD of the E34 TSP. The global quality estimates as well as the local QMEANS are shown in Figure 15C.

The modeling E34 TSP into both the HBD domain and the receptor binding domain (RBD) 3D structures were carried out using the Swiss-Modeler^8^ program **(Arnold et al., 2006)**. The quality of the HBD model was evaluated to show a QMEAN of 0.66 ± 0.05 and a Ramachandran plot analysis as shown in Figure 15B. For the HBD, the molProbity analysis showed a clash score of 3.09, a 96.83% Ramachandran favored structure, 0.00% Ramachandran outliers, 1.66% rotamer outliers and 0 bad bonds out of 2842 bonds. For the local quality estimate, the first 12 amino acids showed QMEAN of less than 0.3, whereas the rest of the structure showed amino acids having a local QMEAN greater than 0.4.

The RBD, as shown in Figure 16A, on the other hand, showed a lower amino acid identity of 29.29% to the exopolysaccharide biosynthesis protein; however, using the Ramachandran plot analysis, it showed molProbity of 2.95, a clash score of 16.92, 81.87% Ramachandran favored structure, 5.50% Ramachandran outliers, 4.72% rotamer outliers and 1 bad bond out of 6777 bonds. A global quality estimate (QMEAN Global) of 0.37 was recorded for the model, with the best Local QMEAN registered between the 150 to 180 amino acid sequence sites, averaging over QMEAN score of 0.55. In general, the modeled RBD structure of the E34 TSP consisted of parallel beta-helices that run orthogonally to one another, showing an overall topology that resembles the RBD of the P22 TSP **(Steinbacher et al., 1997)**.

**Figure 16.**
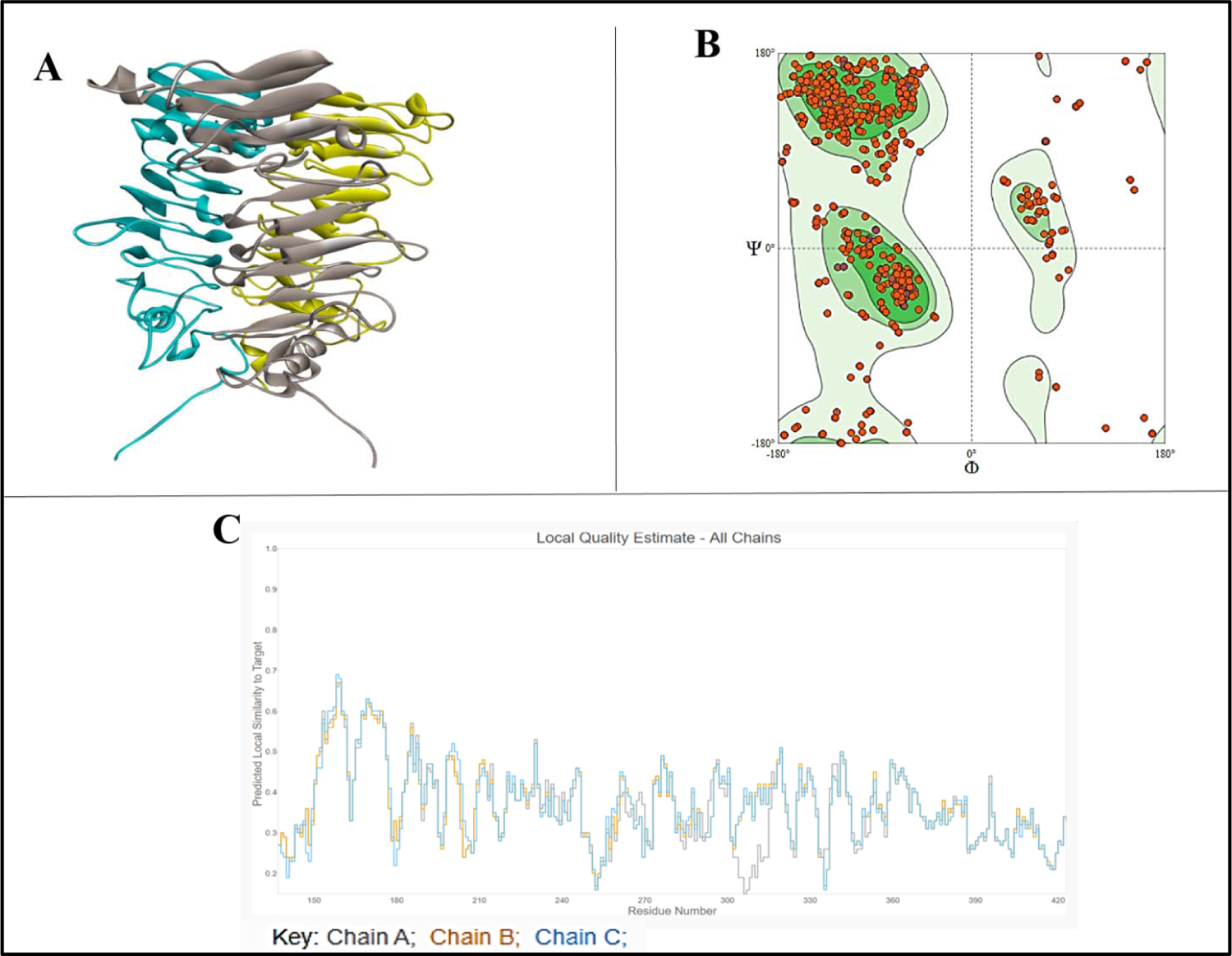
Homology model of the RBD of E34 TSP. Using the Swiss-Modeler, the receptor binding domain (RBD) of the E34 TSP (**Figure 16A**), which consist of the residues from D137 to Y423 amino acids of the E34 TSP were modeled and its stereo-chemical stability evaluated using Ramachandran plot (**Figure 16B**). PDB ID 6TGF.1.A (Irmscher et al., 2021) that shows 29.29% identity was used for the homology modeling of the receptor-binding domain of the E34 TSP. The global quality estimates as well as the local QMEANS are shown in **Figure 16C**.

#### 3.6.4. E34 TSP-S. newington LPS interaction site

The E34 phage is known to interact with the mannoses-rhamnose-galactose repeats of the *S. newington* LPS. The site of this interaction on the E34 TSP is currently unknown. In this study, by computational docking, we computed for the LPS-binding sites of the E34 TSP. This search was initiated by docking a short mannoses-rhamnose-galactose repeat derivative called the mannosyl-rhamnosyl-galactose (PubChem structure CID: 129729227) to the TSP of the phage. Similar to P22 TSP, the receptor-binding site of the E34 TSP was shown to be at the middle domain (RBD) (Figures 17 and 18). As shown in Figure 17D, the 2D depiction of the interaction between the LPS ligand and the E34 TSP receptor indicates that the highest relative free binding energy recorded was between the amino acid positions 250 to 280 of the E34 TSP and it recorded a relative free energy of −6.4 kcal/mol (Figure 18B). Their interaction produced a conventional hydrogen bonding between the ALA250 (bond distance, 2.52Å), SER279 (bond distance, 2.95Å), ASP280 (bond distance, 2.082Å), and a carbon-hydrogen bond at GLY249 (bond distance, 3.29Å).

**Figure 17.**
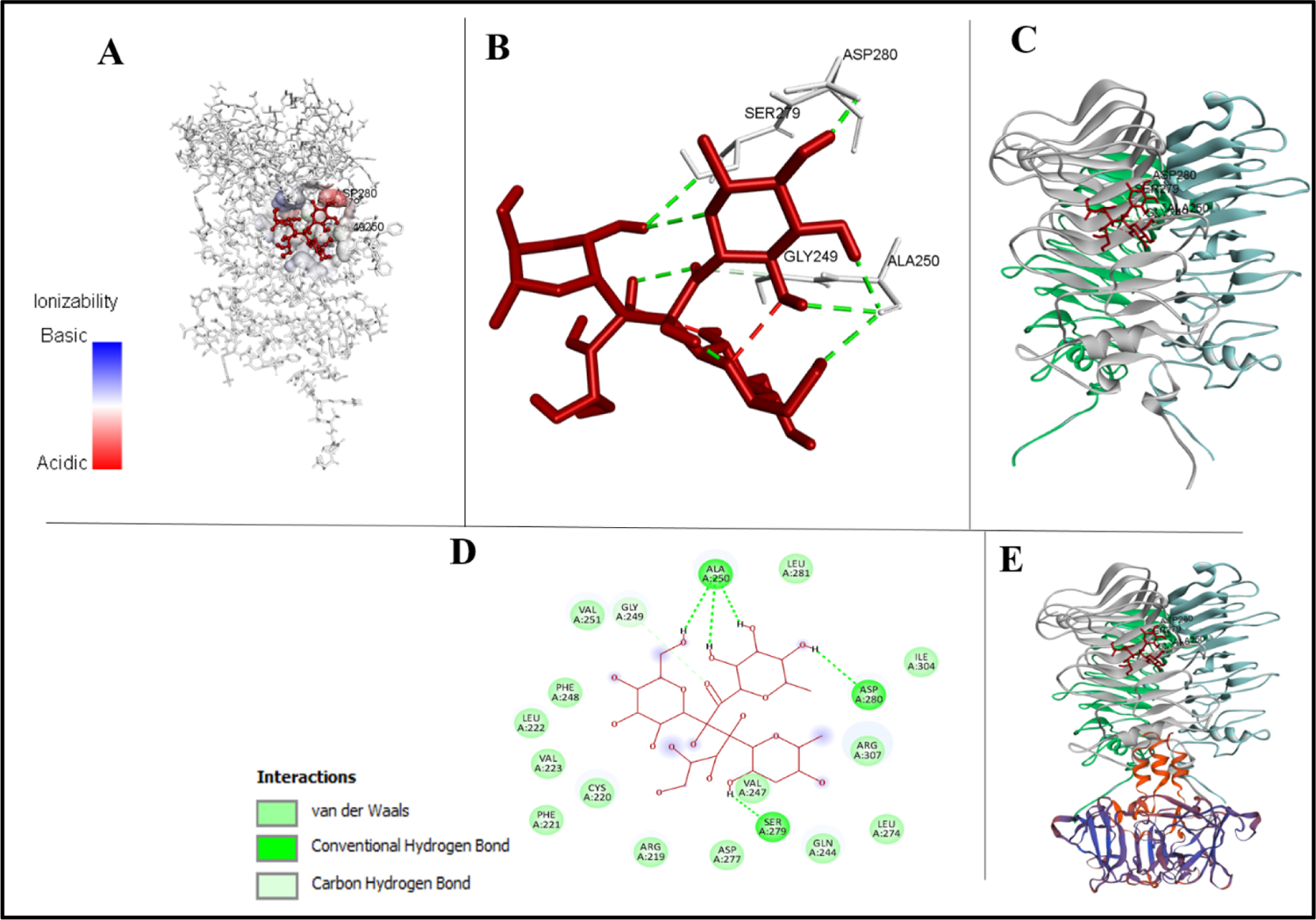
3D and 2D depiction of mannosyl-rhamnosyl-galactose unit binding to RBD of E34 TSP. Interaction between the modeled RBD of E34 TSP and modeled the mannosyl-rhamnosyl-galactose unit (the O-antigen or ligand) after docking with PyRx molecular docking software was analyzed by Biovia Discovery Studio. The RBD depicted in grey sticks whereas the O-antigen subunit is shown in red (**Figure 17A**). It was observed that the highest free energy released was recorded between the binding of the ligand to the SER279-ASP280-GLY249-ALA250 residues (**Figure 17B**). The cartoon depiction of the binding of the ligand to the RBD of E34 TSP (Figure 17C). The 2D demonstration of the binding of the O-antigen to the binding site of the RBD of E34 TSP (**Figure 17D**). **Figure 17E** shows the complete E34 TSP model depicted in cartoon. Structures were visualized with Biovia Discovery Studio and PyMOL (Schrodinger, LLC).

**Figure 18.**
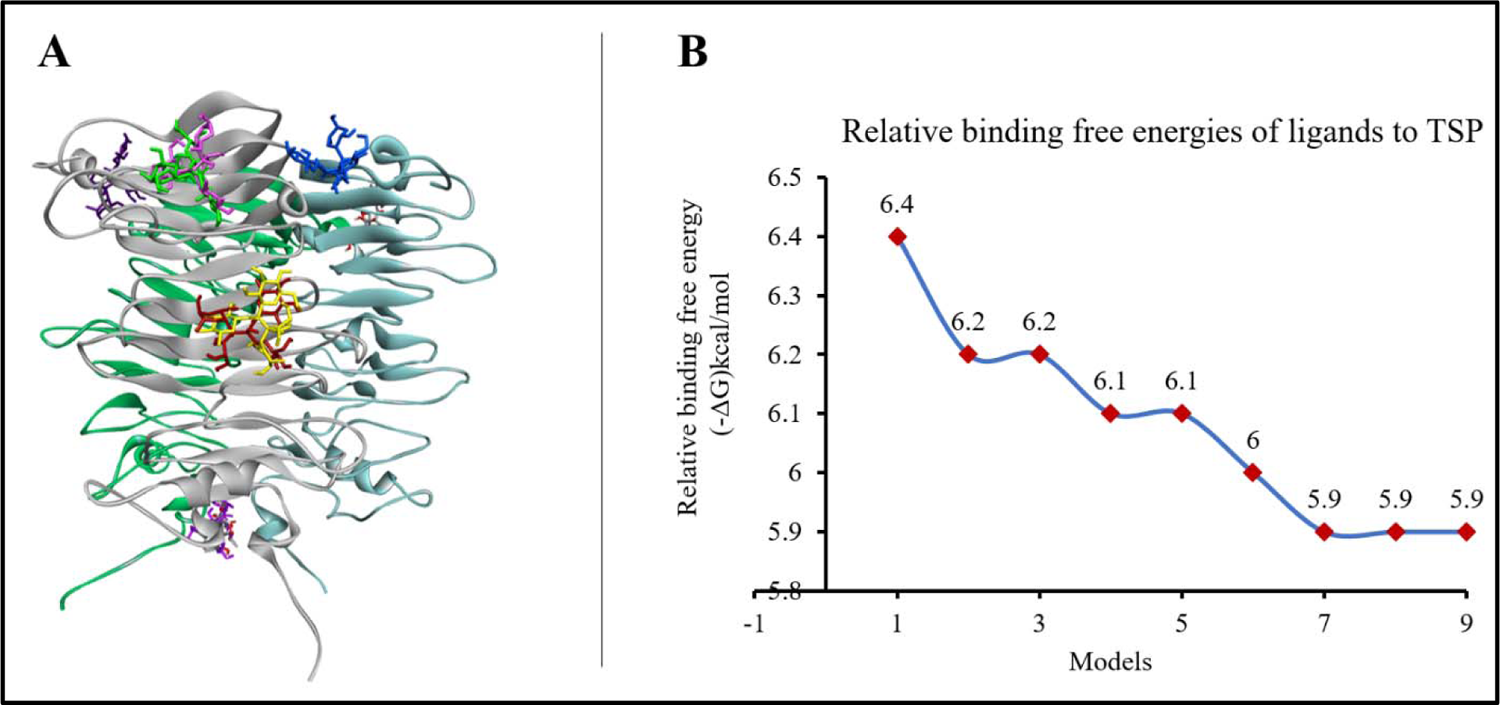
Interaction between the modeled RBD of E34 TSP and the O-antigen ligand. The interaction between the modeled RBD of E34 TSP and the O-antigen ligand showed that the mannosyl-rhamnosyl-galactose unit several different sites of the protein (**Figure 18A**), however the most stable binding occurred at the site SER279-ASP280-GLY249-ALA250 which also recorded the highest free energy (−6.4 kcal/mol) as shown in **Figure 18B**.

##### Vero cell growth in varying concentrations of E34 phage

The Vero cells treated to varying concentrations of the E34 phage showed insignificant difference in growth comparatively, however, relatively higher growth were recorded for cells treated to lower concentration of the phage than those that received larger dosages. As shown in Figure 19, the highest absorbance was recorded for cells treated with the lowest concentrations of E34 phages (2.33 × 10^-4^ µg/ml and 2.33 × 10^-5^ µg/ml µg/ml). An absorbance of 1.83 was recorded for E34 phage treatment at concentration of 2.33 × 10^2^ µg/ml, whereas, 2.29 was recorded for the lowest concentration of E34 phage treatment, thus producing a difference of 0.46 in absorbance between the two E34 phage concentration extremes. The highest absorbance (2.37) however was registered at 2.33 × 10^-4^ µg/ml. The immunofluorescent images (phages showing different multiplicity of infection showed healthy Vero cells attached to the basement of the six-well plates.

**Figure 19.**
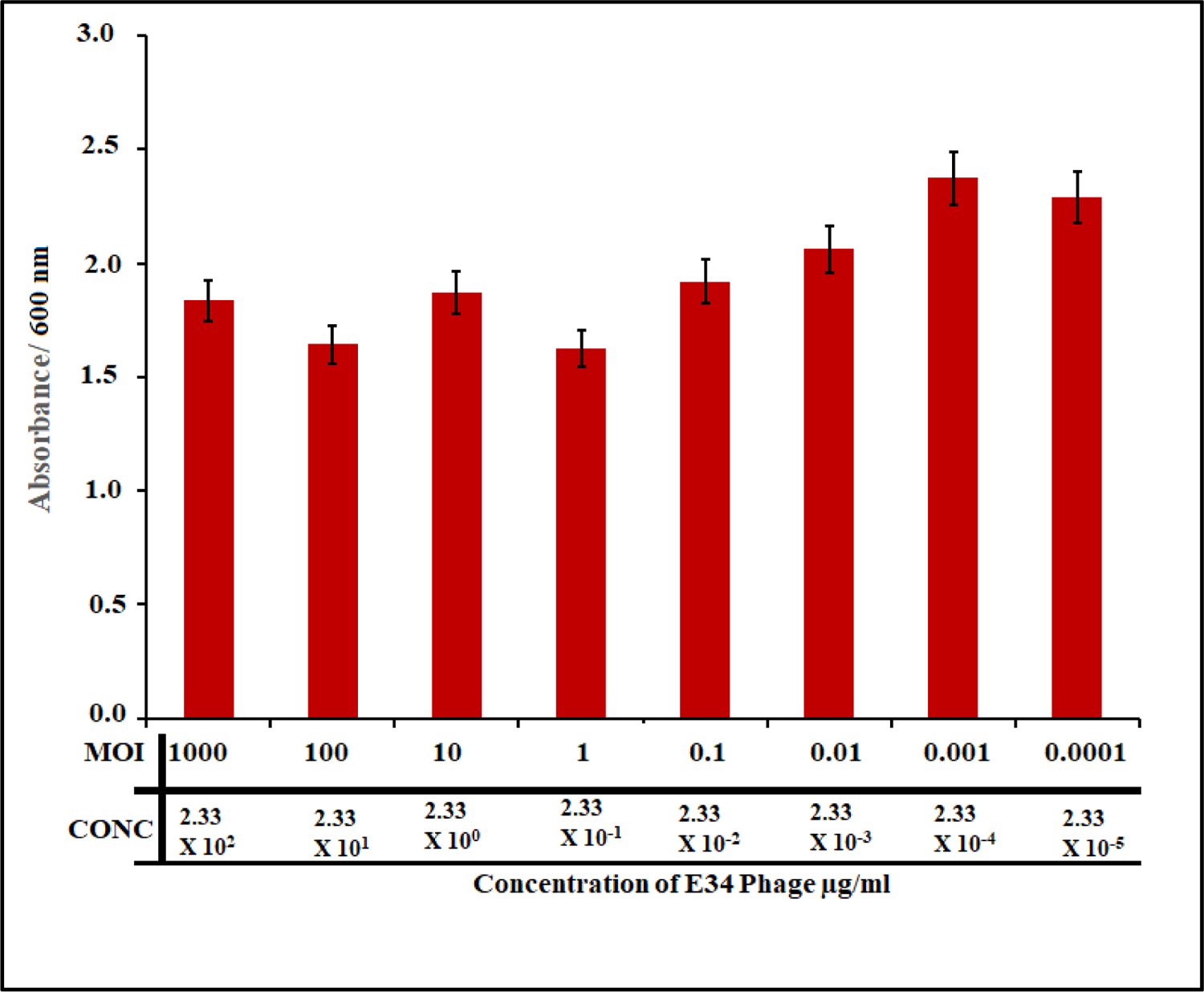
In this experiment, Vero cells were seeded at a density of 1 × 10^5^ into 96 wel plates, then serial dilutions of E34 phages (2.33 × 10^2^ µg/ml to 2.33 × 10^-5^ µg/ml) wer added to cells and incubated for 24 hours. The Vero cells-phage mixture was incubate at 37 °C, 5% CO2 for 24 hours. Wells were then washed twice with 1X PBS to remov dead cells in suspension and fixed with formaldehyde. Fixed cells were the permeabilized using 2% SDS solution and stained with trypan blue, washed twic again, and read at 600 nm in the Cytation 3 plate reader (Biotek, USA). As shown i Figure 19 above, the highest absorbance was recorded for cells treated with the lowes concentrations of E34 phages (2.33 × 10^-5^ µg/ml and 2.33 × 10^-5^ µg/ml µg/ml). An absorbance of 1.83 was recorded for E34 phage treatment at concentration of 2.33 × 10^2^ µg/ml, whereas, 2.29 was recorded for the lowest concentration of E34 phage treatment, thus producing a difference of 0.46 in absorbance between the two E34 phage concentration extremes. The highest absorbance (2.37) however was registered at 2.33 × 10^-4^ µg/ml.

**Figure 20.**
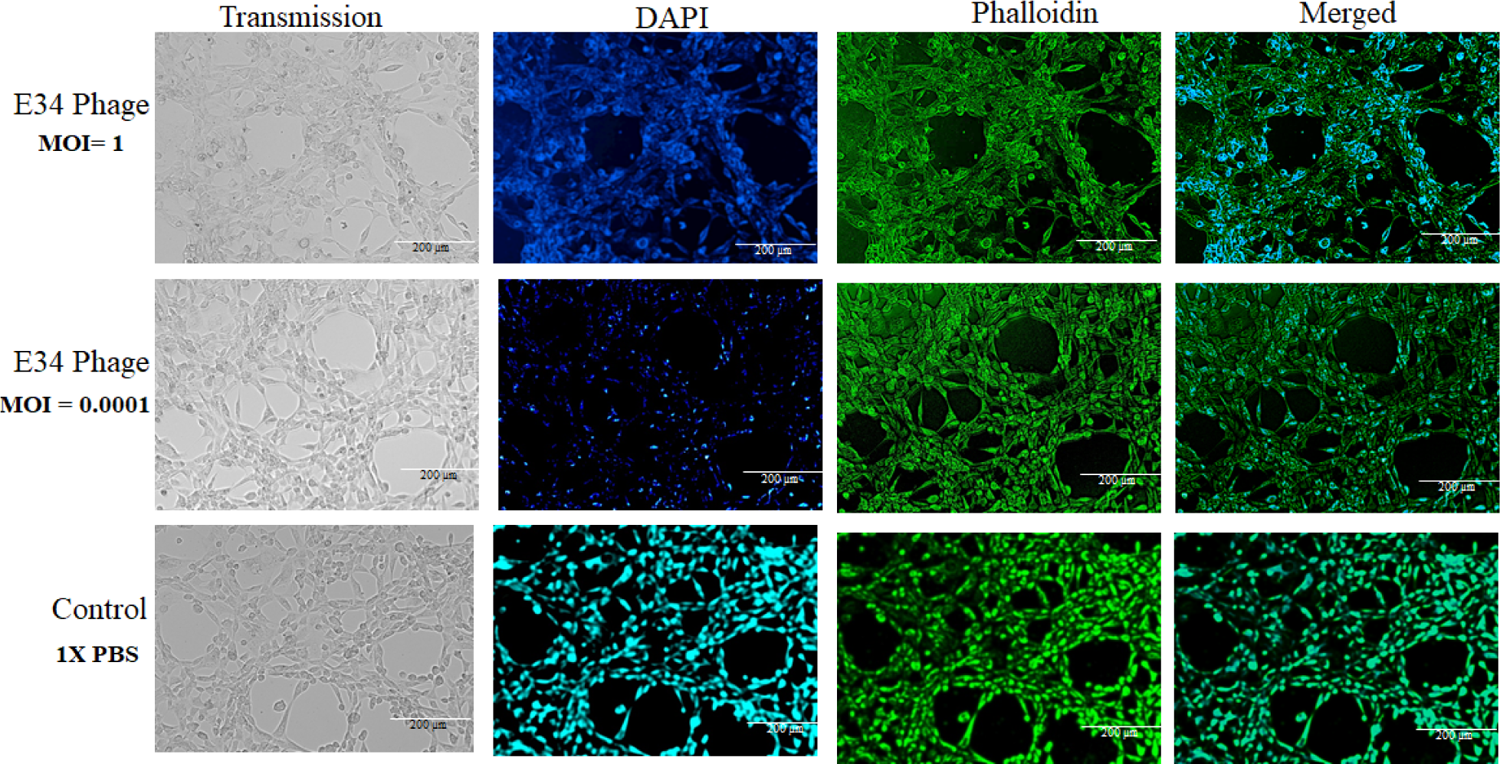
Immunofluorescent images of Vero cells infected with E34 phages are different multiplicity of infection

##### Vero cells are poorly inhibited by E34 phages even at high MOI

To investigate if administration of the phages to the animal cells do have any inhibitory effect of their growth, Vero cells were seeded at a density of 1 × 10^5^ into 96 well plates and followed with the addition of different concentrations of E34 phages. As shown in Figure 21, the highest concentration of phages produced averagely the highest inhibitory effect (mean inhibition of 0.49), whereas the lowest concentration produced the lowest inhibition (mean inhibition of 0.14). The trajectory of the decline in inhibition was fitted to a curve with a slope of −0.0479 and an R^2^ value of 0.9339, which is indicative of high correlation of concentration of phages to Vero cell growth inhibition. In general, although this gentle decline of inhibition is observed across the concentration gradient of the phage treatment, at p-value 0.05, all treatments showed no significant statistical differences at p-value of 0.05. This is also aptly supported by the slope which recorded − 0.0479.

**Figure 21.**
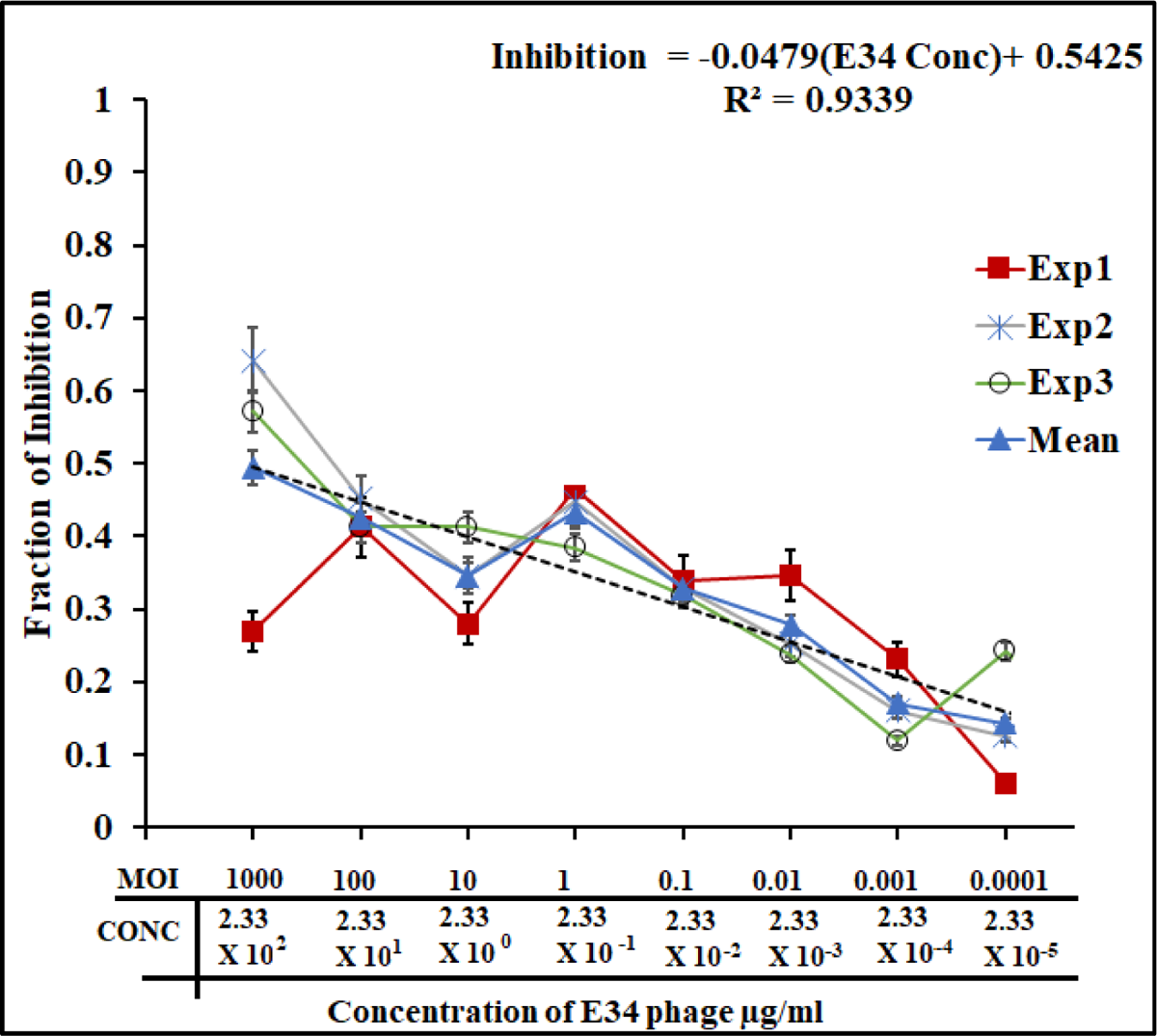
Vero cells are poorly inhibited by E34 phages even at high MOI. In this experiment, Vero cells were seeded at a density of 1 × 10^5^ into 96 well plates an followed with the addition of different concentration of E34 phage and incubated at 3°C at 5% CO2 in Forma Series 3 Water Jacketed CO2 Incubator (Thermo Scientific) for 2 hours. Cells were cultured in DMEM media (ATCC, Manassas, VA) and supplemente with 10% FBS (ATCC, Manassas, VA). Vero cells were treated with E34 phages (experiments, each with 3 replicates) and the ability of the phages to inhibit Vero cell growth were assessed. The highest concentration of phages produced averagely th highest inhibitory effect (mean inhibition of 0.49), whereas the lowest concentratio produced the lowest inhibition (mean inhibition of 0.14). The trajectory of the decline i inhibition was fitted to a curve with a slope of −0.0479 and an R^2^ value of 0.9339, whic is indicative of high correlation of concentration of phages to Vero cell growt inhibition. 1X PBS administration to Vero cells served as the control from whic calculations for inhibition were based on. In general, although this gentle decline o inhibition is observed across the concentration gradient of the phage treatment, at p value 0.05, all treatments showed no significant differences. This is also aptly supporte by the slope which recorded −0.0479.

##### E34 phage protects Vero cells from Salmonella infection

To investigate if E34 phage could protect Vero cells from Salmonella infection, Vero cells were seeded at a density of 1 × 10^5^ into 96 well plates were incubated with S. newington (red line; UC1698 strain), S. newington (BV7004, grey line), and S. typhimurium (BV4012, black line). Approximately, 10^4^ CFU/ml of bacteria cells were used in each treatment. This was followed quickly with the inoculation of varying concentrations of E34 phages (2.33 × 10^2^ µg/ml to 2.33 × 10^-5^ µg/ml). Plague assays were carried out to determine the multiplicity of infection (moi) for each the concentrations to be approximately 1000, 100, 10, 1, 0.1, 0.01, 0.001 and 0.0001. The Vero cells-bacteria-phage mixture was incubated at 37 °C for 24 hours. As shown in **Figure 22**, the phage treatment to Vero cells infected with UC1698 strains produced the highest absorbance in all treatments. At phage concentration of 233 µg/ml, an absorbance of 1.8 is recorded. The highest absorbance of 1.9 was registered at 0.233 µg/ml of phage treatment, this dropped gently to 1.43 at the 2.33 µg/ml × 10^-5^ µg/ml concentration of phage. Cells infected with BV4012 and BV7004 both showed significantly lower absorbance. While the infection with BV7004 recorded 1.37 at 233 µg/ml phage treatment, it continually fell in absorbance to register its lowest of 1.10 at 2.33 × 10^-4^ µg/ml, the overall average absorbance registered for cells infected with BV4012 averaged less than 1.03 in all concentrations. Treatment with 1X PBS media instead of phage produced a very low mean absorbance of 1.16. **Figure 22**), for the UC1698/E34 group, higher concentration of the phage produced higher absorbance because of the lytic nature of the phages that killed most of its bacterial hosts, hence allowing the Vero cells to proliferate. Similar effects are observable in BV7004, however at a lower rate. Both UC1689 and BV7004 are hosts of E34 phage, however, data from our laboratory shows that E34 phage produces slower lytic activity on BV7004 strains as compared to UC1698 (Data not shown), and this slower activity can be observed in this study too, as significantly lower absorbance were registered compared to the UC1698 treatment group. BV4012 is S. typhimurium, which is not E34 phage’s host; thus the presence of the phage does not affect the activity of the bacteria. The BV4012 strains therefore infects the Vero cells without any inhibition from the phage and kills them; and the dead cells are washed off during the washing stage of the experiment; thus lower absorbance values were recorded for all BV4012 treatment.

**Figure 22.**
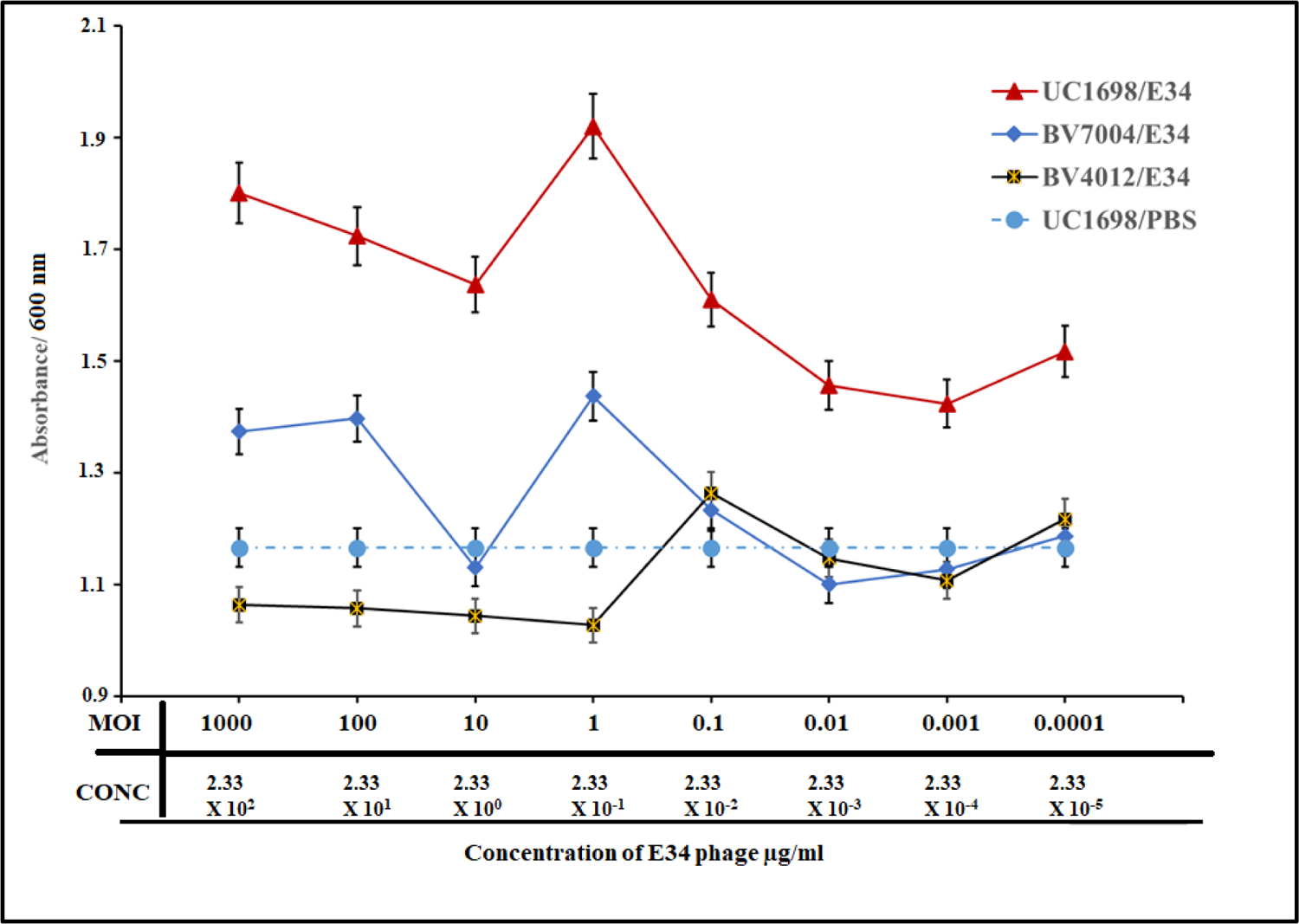
E34 phage protects Vero cells from Salmonella infection The red line representing the incubation of Vero cells with UC1698 (the S. newingto strain that is the host of E34 phage) and treatment to E34 phage. The blue line representing the incubation of Vero cells with BV7004 (the S. newington strain that is th host of E34 phage; slower lytic activity) and treatment to E34 phage. The black line representing the incubation of Vero cells with BV4012 (the S. typhimurium strain that is not the host of E34 phage) and treatment to E34 phage (as a negative control). The broken line representing the incubation of Vero cells with UC1698 (the S. newingto strain that is the host of E34 phage) and treatment to 1X PBS (negative control 2). In this experiment, Vero cells were seeded at a density of 1 × 10^5^ into 96 well plates wer incubated with S. newington (red line; UC1698 strain), S. newington (BV7004, grey line) and S. typhimurium (BV4012, black line). Approximately, 10^4^ CFU/ml of bacteria cell were used in each treatment. This was followed quickly with the inoculation of varying concentration of E34 phages (2.33 × 10^2^ µg/ml to 2.33 × 10^-5^ µg/ml). Plague assays were carried out to determine the multiplicity of infection (moi) for each the concentrations to be approximately 1000, 100, 10, 1, 0.1, 0.01, 0.001 and 0.0001. into the Vero cells-bacteria cultures. The Vero cells-bacteria-phage mixture was incubated at 37 °C for 24 hours. Wells were then washed twice with 1X PBS to remove dead cells in suspension and fixed with formaldehyde. Fixed cells were then permeabilized using 2% SDS solution and stained with trypan blue, washed twice again, and read at 450 nm in the Cytation 3 plate reader (Biotek, USA). As shown in Figure 22, the red line which represents phage treatment to Vero cells infected with UC1698 strains produced the highest absorbance in all treatments. At 233 µg/ml phage treatment, an absorbance of 1.8 is recorded. The highest absorbance of 1.9 was registered at 0.233 µg/ml of phage treatment, this dropped gently to 1.43 at the 2.33 µg/ml × 10^-5^ µg/ml concentration of phage. Cells infected with BV4012 and BV7004 both showed significantly lower absorbance. While infection with BV7004 recorded 1.37 at 233 µg/ml phage treatment, it continually fell in absorbance to register its lowest of 1.10 at 2.33 × 10^-4^ µg/ml, the overall average absorbance registered for cells infected with BV4012 averaged less than 1.03 in all concentrations. Treatment with 1X PBS media instead of phage produced the least absorbance of 1.16 mean absorbance. From the graph, for UC1698/E34 group, higher concentration of the phage produced higher absorbance because of the lytic nature of the phages that killed most of its bacterial hosts, hence allowing the Vero cells to proliferate. Similar effects are observable in BV7004, however at a lower rate. Both UC1689 and BV7004 are hosts of E34 phage, however, data from our laboratory shows that E34 phage produces slower lytic activity on BV7004 strains as compared to UC1698 (Data not shown), and this slower activity can be observed in this study too, as significantly lower absorbance were registered compared to the UC1698 treatment group. BV4012 is S. typhimurium, which is not E34 phage’s host; thus the presence of the phage does not affect the activity of the bacteria. The BV4012 strains therefore infects the Vero cells without any inhibition from the phage and kills them; and the dead cells are washed off during the washing stage of the experiment; thus lower absorbance values were recorded for all BV4012 treatment.

As shown in **Figure 23**, the immunofluorescent images of Vero cells infected with Salmonella newington and counter-treated with E34 phage showed very health phenotypes at lower multiplicity of infection whereas at higher MOIs, Vero cells are seen rounded and with unhealthy features.

**Figure 23.**
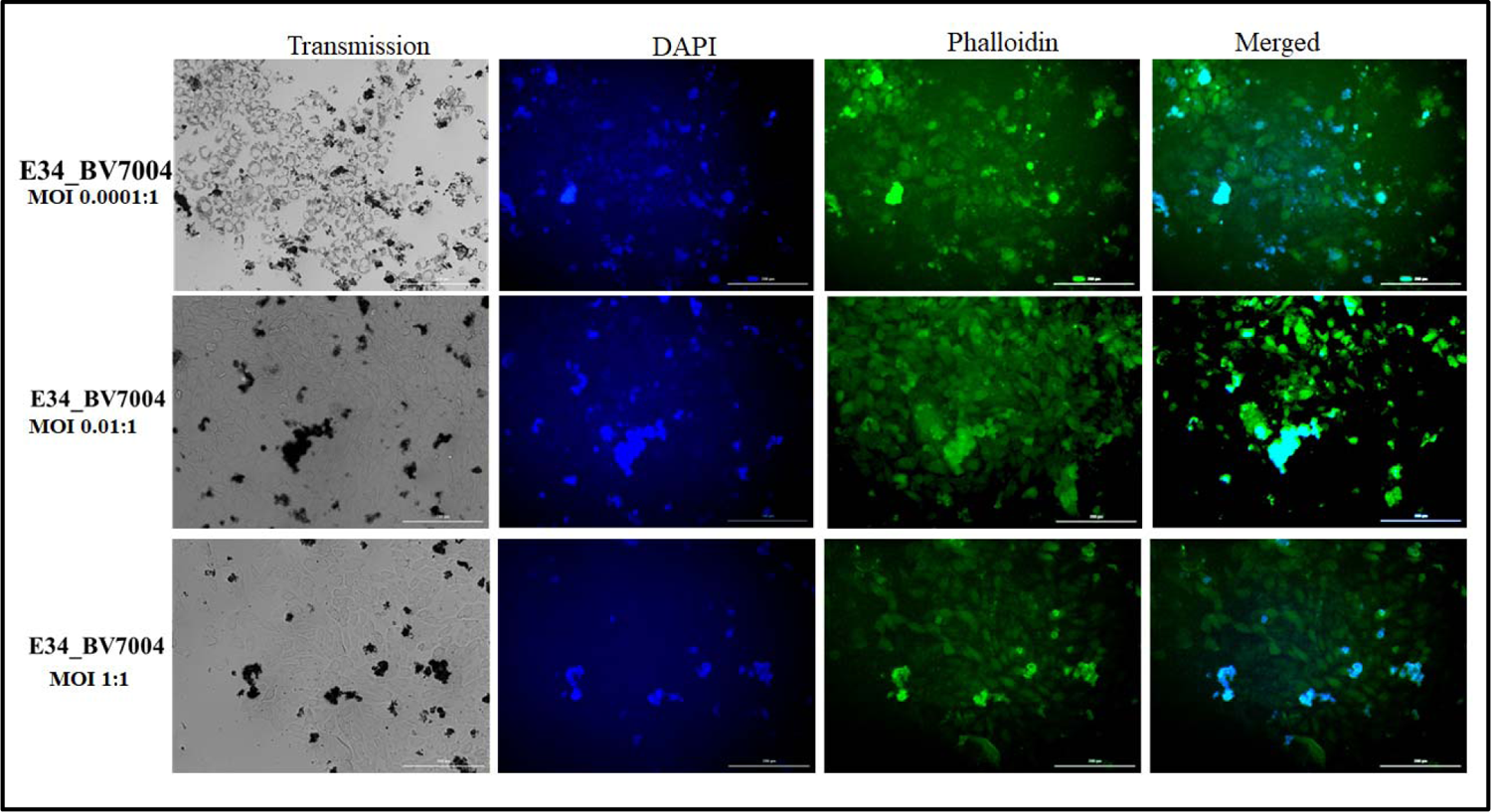
Immunofluorescent images of Vero cells infected with Salmonella newington and counter treated with E34 phage are different MOIs.

In summary, this algorithm evaluated the interaction between the LPS moiety and the E34TSP molecule base on the functional groups on the amino acid, the identity of the amino acid, the surface topology, and the electrostatics occurring between the TSP and the LPS moiety. The relative free binding energy recorded between the interaction between the E34 TSP and mannosyl-rhamnosyl-galactose at this specific site yielded −6.4kcal/mol. Other sites bonded to the LPS moiety recorded lower relative free binding energies, as shown in Figure 18A and 18B. The deficiency of this study, however, lies directly in our choice of molecule used in the study, while mannosyl-rhamnosyl-galactose might be a close derivative of mannose-rhamnose-galactose repeat, it does not exactly feature the exact binding surface, nor the exact electrostatic properties present in the mannose-rhamnose-galactose repeat of the LPS. However, we are confident that this molecule is close enough for a predictive study such as this. Steinbacher et al., 1997 demonstrated that the active site of P22 RBD was the Asp-392, Asp-395, and Glu-359; which acts as catalytic residues **(Steinbacher et al., 1997)**. In this study, our docking results indicate that the endorhamnosidase activity of the E34 TSP might take place between residues 190 and 280 and that the specific binding site of the TSP to the LPS moiety occurred at ALA250, SER279 and ASP280, thus the ASP280 might serve as a nucleophile similar to what is observed in the P22 TSP in which the Asp395 serve as a nucleophile and attacks the anomeric C-atom of O-antigen of the LPS of its host **(Steinbacher et al., 1997).** The active site triad of the serine protease alpha-chymotrypsin is also known to consist of the popular triad HIS-SER-ASP (HIS57, SER195, and ASP102) **(Frigerio et al., 1992)**. In this study, however, the E34 TSP RBD’s active site contained the ALA-SER-ASP triad. The RBD of E34 TSP seems to be a conserved domain, also found in pectate lyases of *Ewinia* and *Bacillus* and the alkaline protease of *Pseudomonas aeruginosa* **(Yoder et al., 1993; Pickersgill et al., 1994; Yoder and Jurnak, 1995).**

## Statistical analyses

For all statistical data, values were derived from multiple measurements (from replicate of 3 or 5 experiments) and averaged; the standard deviations were evaluated using P values of Student’sst (one tailed, two samples of unequal variance, significance level α = 0.05).

## 4. Discussion

E34 phage, a podovirus infects *S. newington*, and the TSP of this phage is responsible for the attachment of the phage particle to the LPS of the cell and subsequent hydrolysis and anchoring of the same to the membrane of the cell to initiate the phage’s injection of its DNA into the cell. In this study, the E34 TSP, the protein derived from the gene gp19 coding sequence, was cloned and expressed under the control of the T7 promoter in the pET30a-LIC vector, which adds 43 amino acid N-terminally to the protein. This fusion protein, EE34 TSP, contained the His-tag suited for affinity chromatography.

Validation of the cloned insert was achieved via three methods: 1) PCR reaction using two primers that amplified exactly the tailspike gene, gp19 in the clone, and verification by agarose gel electrophoresis where the putative clones (recombinant pET30a-E34 TSP, 7.2 kB) were the size of the vector (5.4 kB) and the inserted tail gene fragment (1.8kB). 2) The 210kDa trimeric protein (196kDa TSP + 14kDa (three 43aa fusion peptide) was cleaved successfully with rEK enzyme, which cleaves at a site (DDDDK) between the His-tag region and the E34 TSP to produce the wildtype protein E34 TSP and the His-tag fragment (Fig. 2). 3) The His tag was verified by monoclonal antibody to the His-tag amino acid sequence (data not shown). Ni-NTA column was used for affinity purification via FPLC (Fig. 1).

The P22 TSP structure had previously been shown to contain a structure that are stable to heat (Tm of 88.3 °C), proteases, and to detergents **(Williams et al., 2019)**. Therefore, it was of interest to determine if the EE34 TSP was part of this class of proteins. Manning and Colon demonstrated that rigid protein structures containing oligomeric beta-sheets are implicated for the physical basis for kinetic stability, SDS resistance, as well as protease resistance of proteins with such constitutions **(Manning and Colón, 2004)**. We have demonstrated via 1) online BETAWRAP program that EE34 TSP contains several oligomeric beta-sheets (data not shown), 2) Swiss-models (Fig. 16A) shows that our protein consists of extensive beta structures; additionally, Salgado et al. 2004, predicted structural similarities between E34 TSP and P22 TSP **(Salgado et al., 2004)**, the P22 TSP consists of mainly a high percentage of beta structures **(Steinbacher et al., 1994).**

In this study, we subjected EE34 TSP to varying concentrations of SDS and analyzed the kinetics of the protein via native polyacrylamide gel electrophoresis. Our comparator protein, the P22 TSP produced a single discrete band in all concentrations of SDS, indicating stability. The EE34 TSP produced a blurry non-discreet band at 0 % SDS concentration which we propose to represent an ensemble of isomeric forms of the EE34TSP in equilibrium under the non-denaturing buffer solution, but when SDS was added to the samples, these isomeric species were converted into four main groupings.

Increasing concentration of SDS also seemed to chase the band 2 and 3 species into band 1, and 4 because the densities of these bands (band 1 and band 4) increased with increasing SDS concentration with a concomitant decrease in the densities of bands 2 and 3. We hypothesized that isomers in bands 2 and 3 are chased into bands 1 and 4 as SDS concentrations climb above the critical micellar concentration of 0.2 % SDS to account for such transformation. In our proposition, the numerous isomers of EE34 TSP are driven into the four main band categories via their interaction with SDS molecules. Thus, these interactions can generate the free energies necessary for isomers high up the energy funnel to reach their activation state enabling them to cross the energy barrier and fall to the native state or to isomeric forms that are much closer to the native state. Thus, we infer those energies generated from TSP-SDS micellar interactions converted the ubiquitous ensembles of EE34 TSP into only the four groups with the four groups reconverted into only two as concentrations of SDS increased to exceed the critical micellar concentrations. In summary, this interaction between SDS and EE34 TSPs might indicate a structural reshuffling for which less stable EE34 TSP species are converted to a more stable form.

The EE34 TSP has a 43-amino-acids fragment at the N-terminus of the E34 TSP, it was of interest to determine if this addition might affect the structure of the EE34 TSP. Use of the PONDR program **(Xue et al., 2010)**, allowed the prediction of the structural lability of the 43 amino acids fusion peptide placed at the N-terminal end of EE34 TSP. This program indicated that the fusion peptide was predicted to be an intrinsically disordered region (IDR) end of the protein. IDPs with unstructured domains, sometimes under slightly altered environments, can additively corroborate to attain harmful and deleterious forms of proteins that are suspects in most common misfolding diseases **(Oldfield et al., 2008)**. Understanding the behavior of our protein under the varying concentration of SDS may provide a novel method that can be used to study IDPs of major clinical importance, such as the p53 and the proto-oncogene, c-myc.

The thermal stability characteristics of EE34 TSP, presented in this study reveals subtle similarities to the well-studied P22 TSP, as it is not surprising that both proteins are homotrimers, consist largely of beta structures, are both involved in endorhamnosidase activity, and have been proven to possess over 70% identity in their N-terminus head binding domain as determined by amino acid sequence comparisons. With both proteins stable at higher temperatures, our thermal stability data strongly indicate that the P22 TSP is more heat stable than EE34 TSP, since, at 90 °C, P22 TSP still existed at least 50% as trimeric species, while at the same temperature, all species of EE34 TSP had fully denatured. This agrees with several other studies on P22 TSP **(Salgado et al., 2004; Zayas and Villafane, 2007; Chen and King, 1991; Casjens and Thuman-Commike, 2012; Williams et al., 2019; Fuchs et al., 1991)** and the initial characterization and prediction made by Zayas and Villafane of E34TSP **(Zayas and Villafane, 2007)**. Heating at 70 °C only minimally affected the EE34 TSP, while the P22 TSP was unaffected. At higher temperatures, the EE34 TSP showed unfolding, with a majority of EE34 TSP species in their monomeric state while P22 TSPs remained unaffected. Heating P22 TSP at 80 °C did not affect the protein even after an hour of incubation. The unfolding pathway observed for EE34 TSP did not follow an initial intermediate before subsequent complete unfolding into monomers as observed in most thermal unfolding kinetic studies in P22 TSP **(Chen and King, 1991; Danner et al., 1993; Fuchs et al., 1991; Goldenberg and King, 1982)**, but fell from trimeric native species into monomers. The P22 TSP is much more stable to temperatures from 65°C – 90 °C **(Chen and King, 1991; Danner et al., 1993; Goldenberg and King, 1982)**. It is important to note that even though P22 TSP possess higher thermal stability than the EE34 TSP, we could still infer a high degree of similarities in their thermal characteristics in the fact that even a single point mutation in P22 TSP has been demonstrated to decrease or increase the denaturation rate constants several folds respectively **(Miller et al., 1998; Goldenberg and King, 1982),** hence, such huge dissimilarities in sequence identity and yet both proteins possessing such marginal difference in thermal stability points to their identity in overall structural topologies instead of amino acid sequence identity.

In this work, we also propose that the observed plasticity of conformations in this protein is caused by the unstructured 43 amino acid fusion peptide; and this aberrant mobility phenomenon has been identified with the other intrinsically disordered proteins **(Iakoucheva et al., 2001)**. The secondary structure of these 43 amino acid fusion peptides is predicted to be composed entirely of an exposed and solvent-accessible strand, with a 100% disorder that lacks structure. The lack of structure of the fusion peptide gives it the unlimited possibilities of interaction **(Apellániz et al., 2014; Oldfield et al., 2008; Andresen et al., 2012)** with the more structured components of the protein or with similarly unstructured regions in both intra- and inter-monomeric interactions during the folding process **(Oldfield et al., 2008)**. For instance, the fusion peptide strand can interact with the slightly solvent-accessible regions of the more structured N-terminus in the same monomer. In another scenario, the same monomer can interact with the exposed neck region, which links to the midsection of the monomer via polar electrostatic interactions, or in a trimeric conformation, the fusion peptide could interact with the two other fusion peptides or with the structured N-terminus which can lead to the production of distinct but closely related conformational species with varying energetics. In the presence of SDS or other denaturants, most of these exposed sites are bonded to and occupied by the denaturing molecules leaving limited sites available for the fusion peptide strand to interact with. We propose that the limitation in interaction sites for the fusion peptide forces the protein into a limited number of conformations as seen in SDS-PAGE analysis of the EE34 TSP in which at lower concentrations of SDS four bands are observed. However, as SDS concentration increases, the mechanism of interaction between SDS and TSP changes **(Varela et al., 2018; Klaus and David, 1971; Abani, 2009)**, from the two forms of interactions (which is both hydrophilic and hydrophobic interactions) at lower SDS concentrations to mainly a single form of interaction which is hydrophobic; and this change further limits the number of isoforms produced, hence the limited number of bands (e.g., 2 main bands observed at higher SDS concentrations).

The half-life of EE34 TSP in a solution containing 0.2% SDS set at different temperatures was well illustrated via the unfolding midpoints observed at the set temperatures, a half-life of 125 mins at 50 °C, followed by a dropped to 15 mins in 70 °C, then a further drop to 5 mins at 80 °C, and finally 29 sec at 90 °C. These data demonstrate the cooperative unfolding mechanism of both SDS and heat on the EE34 TSP. Similar studies have shown that at 65°C, the P22 TSP, which is noted for its thermostability, was fully denatured when treated in an SDS-heat combo within 30 mins **(Chen and King, 1991)**. At the temperatures tested, the EE34 TSP was more heat sensitive than the P22 TSP (Tm of 88.3 °C) **(Sturtevant et al., 1989)**.

In analyzing if EE34 TSP interferes in P22 H-P22 TSP interaction, it is clear from the data produced that binding of EE34 TSP to P22 H competitively excludes P22 TSPs from binding to the P22 H since no plagues were observed at higher concentrations of EE34 TSP but could be observed at the lower concentrations of EE34 TSP. This occurred because all available binding sites in the P22 H for the P22 TSP had been occupied by EE34 TSP; this way, EE34 TSP interferes in P22 TSP-P22 H binding. This blocking of P22 TSP from binding to its P22 H by the bound EE34 TSP interferes in the assembling process of the P22 phage into a complete infective phage. This explains the presence of the plagues at the last two spots of the P22H + EE34 TSP, P22 TSP lane (Fig. 13). This is confirmed in Fig.14, for which at high concentration of EE34 TSP, a few plaques were recorded, whereas at lower concentrations of EE34 TSP high number of plagues were recorded. The ability of ELJ34 TSP to interfere in the assembly of the P22 phage indicates structural similarity in the N-terminal domains of both proteins. It also points to the fact that the 43 amino acid fragment is labile and unstructured and did not affect the overall structure 34 protein.

As demonstrated by Steinbacher et al., 1997, the binding and also catalytic residues of P22 RBD is the Asp-392, Asp-395, and Glu-359 **(Steinbacher et al., 1997)**. Our virtual screening indicates that the E34 TSP hydrolase active site lie between residues 190 and 280 and that the specific binding site of the TSP to the LPS moiety occurred at ALA250, SER279 and ASP280. We propose that the ASP280 might serve as a nucleophile similar to what is observed in the P22 TSP in which the Asp395 serve as a nucleophile and attacks the anomeric C-atom of O-antigen of the LPS of its host **(Steinbacher et al., 1997)**. Also, the active site triad of alpha-chymotrypsin is HIS57, SER195, and ASP102 **(Frigerio et al., 1992)** our hydrolase with catalytic property such as this enzyme seems to assortment of residues at its active site (the ALA-SER-ASP triad).

The cytotoxicity assessment of E34 phage to Vero cells indicates that these phages do not present major harm to animal cell lines especially at lower concentrations. These findings suggest that E34 phage can be formulated into a tablets/shrubs for *in vivo* administration without causing serious harm to the study animal. In general, we used Vero cells as *in vitro* model to assess the safety and efficacy of E34 phage treatment of *Salmonella* infection. Similar methodology from other researchers have demonstrated similar findings pertaining the safety of some phages in phage therapy **(Shan et al., 2018)**. Nonetheless, this work shows many limitations; first, a monolayer of Vero cells cannot truly recapitulate the complex microenvironment of most tissues (e.g., the lining of the gut where most infections may occur), secondly, cell viability and membrane integrity measurements alone cannot offer full insight into the cellular and biological signaling mechanism that can be induced by the presence of a bacteriophage in a living tissue. Here, we have not shown also any specific target receptor(s), that might be implicated or otherwise in phage-tissue interaction. Notwithstanding these major limitations, this work serve to elucidate the biocompatibility of E34 phage usage in phage therapy.

In summary, these studies provide the unique characteristics of the ELJ 4 TSP, contrasts with the model P22 TSP and highlighted its activity. Additionally, we discovered by in silico analysis the catalytic site of the E34 TSP. This characterization sheds light on the biology of the LJ Bacteria and phages relationship enhances their rapid coadaptation that allows highly dynamic coevolution **(Koskella and Brockhurst, 2014)** an in-depth understanding of the biology of the phages, especially their tailspikes could help develop methods to monitor, treat, or prevent pathogenic bacteria such as *Salmonella*, more so, the understanding of their biology offer industrially applicability, for instance, the development of biosensors, based on the phage tailspike’s properties and specificities to the LPS its host. Finally, testing E34 phage ability to protect Vero cells from Salmonella infection shows highly encouraging results, implying that E34 phage can be used in therapeutic/preventive medicine. Currently, our lab is investigating the bacteriostatic property of this expressed protein against *S. newington*, as well as the protein’s immune-regulatory character on human mesenchymal stem cell lines.

## Acknowledgments

We thank the Department of Biological Sciences, Program in Microbiology in the College of Science, Technology Engineering and Mathematics (C-STEM) Alabama State University for support.

## Ethical Approval

This article does not contain any studies with human participants or animals performed by any of the authors.

## Footnotes

1 www.cdc.gov/ncezid/dhqp

2 www.cdc.gov/ncezid/dhqp

3 http://www.ncbi.nlm.nih.gov/protein

4 http://swiss-model.expasy.org

5 http://mordred.bioc.cam.ac.uk

6 https://pubchem.ncbi.nlm.nih.gov

7 https://www.ncbi.nlm.nih.gov

8 http://swiss-model.expasy.org

